# Dual-encoder contrastive learning accelerates enzyme discovery

**DOI:** 10.1101/2025.08.21.671639

**Authors:** Jason W. Rocks, Dat P. Truong, Dmitrij Rappoport, Samuel Maddrell-Mander, Daniel A. Martin-Alarcon, Toni M. Lee, Steven Crossan, Joshua E. Goldford

## Abstract

The ability to engineer enzymes for desired reactions is a cornerstone of modern biotechnology, yet identifying suitable starting proteins remains a critical bottleneck. Although contrastive learning offers a compelling computational approach for enzyme discovery, these models have yet to be implemented at scale or proven effective in real-world experimental settings. Here, we present *Horizyn-1*, a computationally efficient deep learning framework that enables large-scale reaction-to-enzyme recommendation validated through comprehensive experimental testing. Leveraging a combination of reaction fingerprints and protein language models, we trained *Horizyn-1* on millions of reaction-enzyme pairs to achieve state-of-the-art performance, recovering an enzyme with correct activity within the top 100 hits for over 75% of test reactions. We experimentally validate *Horizyn-1* across three enzyme discovery scenarios: identifying enzymes for orphan reactions, predicting enzyme promiscuity for both characterized and uncharacterized enzymes, and discovering enzymes for non-natural biochemical reactions including lysine-driven transaminations that enable efficient synthesis of non-canonical amino acids. On underrepresented reaction classes, we find that fine-tuning with fewer than 10 additional reactions can dramatically improve performance. Furthermore, a logarithmic scaling of model performance with training dataset size suggests continued improvement with larger and more diverse reaction datasets. *Horizyn-1* addresses the critical bottleneck of sourcing initial enzymes for optimization campaigns, enabling efficient and scalable *in silico* screening for enzymes with desired activities and promising to accelerate future efforts in biocatalysis and metabolic engineering.

## INTRODUCTION

The emerging field of generative models for enzyme design holds immense promise for creating novel biocatalysts [5, 29]. However, the performance and scope of these models are critically dependent on the quality and breadth of the training data, which is dominated by known, wild-type enzymes [27]. Consequently, future advancements in this area require robust tools for the discovery and annotation of enzymes exhibiting novel reactivity [27]. Advances in highthroughput metagenomic sequencing have yielded an explosion in non-redundant protein sequences throughout the last two decades, many of which contain domains of uncharacterized enzymatic function [11]. Identifying previously undescribed catalytic activities within these naturally occurring enzymes presents a grand challenge with significant implications for disciplines as diverse as oncology [35] and microbial ecology [39].

Historically, protein function annotation tools have relied on conventional bioinformatics and sequence-similarity based approaches [1, 6, 13, 33, 48, 52, 56]. More recently, advances in deep learning have enabled models to learn complex sequence-function relationships from large datasets, leading to accurate predictions for functional descriptors such as EC numbers, GO terms, or enzyme-substrate interactions [10, 26, 44, 55]. However, all these approaches are fundamentally limited by their reliance on indirect or incomplete descriptions of catalytic activity instead of direct representations of the biochemical reactions themselves. The BridgIT algorithm, a notable exception, uses similarity between custom reaction fingerprints to identify plausible enzymes for a query biochemical reaction, but lacks any machine learning capabilities [18].

Recent advances in protein language models demonstrate that scaling to larger and more diverse datasets consistently improves performance [30, 34, 41]. These large-scale models have begun to enable practical applications in protein engineering and design [4, 19, 43]. While protein language models excel at capturing sequence features, connecting these sequences to chemical transformations requires a multimodal approach. Dual-encoder contrastive learning is uniquely suited for this task, having proven effective at aligning elements from distinct domains like images and text [40]. Recent studies have utilized this approach to associate reactions and enzymes directly [24, 31, 36, 54], offering a method to screen large sequence libraries for novel or hypothetical activities. Despite promising results on machine learning benchmarks, current implementations have been limited to small training datasets and have not undergone experimental validation. Here, we present a framework that leverages large-scale training and is validated through comprehensive experimental testing.

## RESULTS

### Scalable enzyme annotation and discovery with joint reaction and enzyme embedding

To explore the utility of contrastive models to guide experimental enzyme discovery and annotation, we developed *Horizyn-1* (Fig. 1, see Methods). This model adapts the contrastive learning framework popularized by the CLIP image-to-text model to learn associations between reactions and protein sequences (Fig. 1A) [40]. Input reactions and proteins are converted to vector representations and then projected into a common high-dimensional embedding space using separate multilayer perceptron (MLP) encoders. Within this space, the model is trained to minimize the distance between known reaction-enzyme pairs (positives) while maximizing distances for unrelated pairs (negatives). This process yields a joint embedding space where proximity correlates with the likelihood a protein catalyzes a given reaction, enabling enzymes or reactions to be ranked in order of relevance to a particular query reaction or enzyme (Fig. 1B–C). *Horizyn-1* computes scores via the dot product (cosine similarity, ranging between −1 and 1) between the embedding vectors of a given reaction-enzyme pair.

**FIG. 1.**
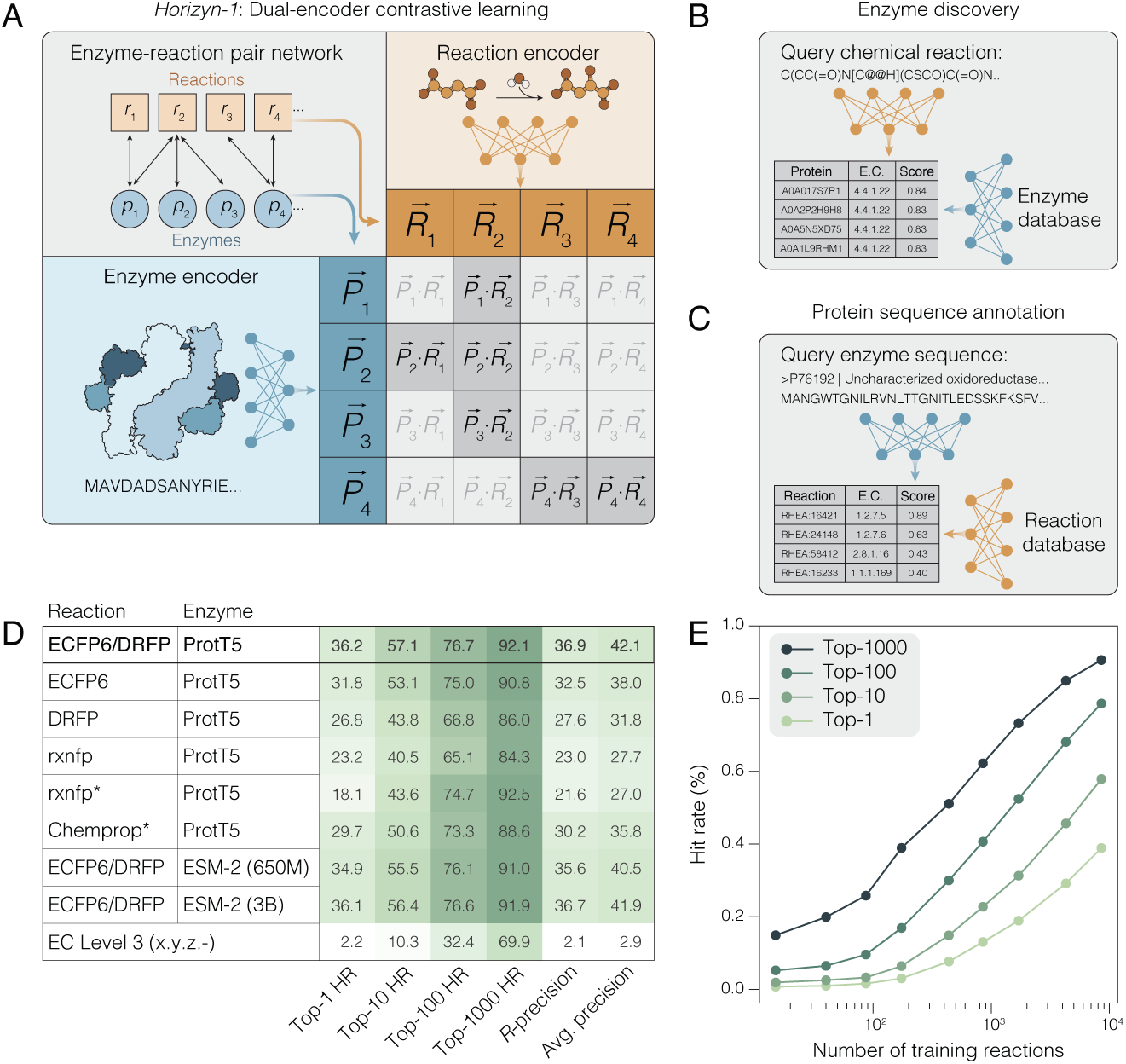
*Horizyn-1* model architecture, applications, and performance. **(A)** Schematic of the contrastive learning framework. Reactions and protein sequences are independently encoded into vector representations and projected into a shared embedding space. The model is trained to minimize the distance between known reaction-enzyme pairs (positives) and maximize the distance between unrelated pairs (negatives). **(B)** Enzyme discovery workflow: Using a query reaction, candidate enzymes are ranked by their proximity in the embedding space, where closer proximity correlates with a higher predicted likelihood of catalysis. The output is a table of enzymes ranked by their score. Enzyme Commission (EC) numbers are also shown for reference. **(C)** Protein sequence annotation workflow: The reverse application, where a query enzyme is used to identify and rank potential reactions it may catalyze. **(D)** Heatmap comparing retrieval metrics, including top-*k* hit rates (HR, *k* = 1, 10, 100, 1000), *R*-Precision, and average precision, for different combinations of reaction and protein representations. The highest-performing model (top row) uses a combination of structural and differential fingerprints (ECFP6/DRFP) with static embeddings from the ProtT5 protein language model. * Denotes cases where reaction embedding models were fine-tuned (rxnfp) [46] or learned (Chemprop) [20] during training. **(E)** Data-scaling analysis showing a logarithmic relationship between model performance (top-100 hit rate) and the number of training reactions. This suggests that performance can be further improved with larger datasets.

For model development, we constructed a training dataset of 294,166 reaction-enzyme pairs derived from 11,961 biochemical reactions in the Rhea database [3], with 216,132 corresponding enzyme sequences from Swiss-Prot [51]. We constructed a representative test set of 1,012 reactions by splitting the data based on reaction similarity, calculated using classical fingerprinting techniques (see Methods). To evaluate model performance, we measured how highly the model ranked enzymes known to catalyze each test reaction relative to all other enzymes in the dataset. We report a variety of standard retrieval metrics (Fig. 1D, Fig. S1, Table S1, Table S2), but focus primarily on top-*k* hit rates, measuring the percentage of reactions in the test set where at least one known enzymatic catalyst was successfully recovered within the top-*k* ranked sequences. Top-*k* metrics are especially informative because they map directly to experimental design: the top-*k* predicted sequences correspond to the number of variants synthesized and screened in a typical enzyme discovery campaign (see Methods).

While we considered a wide variety of different model architectures and training strategies, we found that the choice of strategy for creating reaction and protein representations were most impactful. Our most successful model architecture combined a hybrid reaction representation—using extended-connectivity fingerprints (ECFP6, also known as Morgan fingerprints) that combine the structures of the reactants and products (structural fingerprints) [42, 45] and Differential Reaction Fingerprints (DRFP) that encode the bonding changes (differential fingerprints) [38]—with pre-computed protein sequence embeddings from the 3-billion-parameter ProtT5 protein language model [14]. This combination achieved strong performance, identifying at least one known enzymatic catalyst within the top 100 recommendations for 76.7% of test reactions (Fig. 1D). In contrast, a baseline approach using EC numbers up to the third digit to rank sequences—essentially using detailed knowledge of the reaction class—achieved a top-100 hit rate of only 27.5%. Notably, we found that combining our model with partial knowledge of the EC class could boost top-*k* retrieval metrics by approximately 5–10% (see SI Appendix and Table S3). Substituting the reaction representation with approaches based on transformers (rxnfp) [46] or graph neural networks (Chemprop) [20] generally led to lower scores, with significantly reduced top-1 and top-10 hit rates (Fig. 1D). Similarly, protein embeddings generated with ESM-2 (650M and 3B) [30] resulted in slightly reduced performance, especially for the smaller model (Fig. 1D). Furthermore, neither fine-tuning ESM-2 (650M) nor modifying the architecture to incorporate 3D protein structures offered substantial performance improvements over static embeddings despite significantly increased computational costs (see SI Appendix and Fig. S2).

Our training strategy employed in-batch negative sampling [40], where all mismatched protein-reaction pairs within a mini-batch are treated as negatives. We found that this approach outperformed the use of hard negatives constructed from shared EC classes [55], while also demonstrating superior computational efficiency (see SI Appendix, Table S4). However, this strategy inherently risks treating missing annotations—valid enzyme-reaction pairs that are not yet recorded in the database—as negative examples, potentially causing the model to overfit to the incompleteness of the training data. To assess this, we simulated the effects of missing data by artificially leaving out known positive pairs during training. We found that the model retained strong predictive performance on these missing pairs, indicating robustness against unannotated promiscuous activities (see SI Appendix, Fig. S3).

We directly compared the model architecture and training procedure of our best model with those of CLIPZyme [36], an influential contrastive model for reaction-enzyme association. We found that our model moderately outperformed CLIPZyme, likely due to the use of a larger protein language model (ProtT5 at 3 billion vs. ESM-2 at 650 million parameters) and fingerprint-based reaction representations rather than a learned graph neural network implementation based on pseudo-transition states (see Table S4, Discussion).

To investigate the relationship between training dataset size and model performance, we performed data-scaling experiments by varying the number of training reactions (clustered on reaction fingerprint similarity, see Methods). We found that training with only 10% of the data yielded a model with a top-100 hit rate exceeding 40%. We observed logarithmic scaling of model performance with the number of training reactions, indicating that an approximate tenfold increase in reactions could yield a model with a top-100 hit rate approaching 100% (Fig. 1E). These results suggest that further scaling of the model with a substantially larger—and diverse—set of reactions and protein-reaction pairs could yield a model with enough predictive power to construct high-quality panels of enzymes to guide enzyme discovery campaigns with high confidence.

As model performance improves significantly with both the number and diversity of reactions, we assembled a large dataset of reaction-enzyme pairs by combining multiple databases (totaling ∼31,000 reactions, ∼27 million protein sequences, and ∼34 million reaction-enzyme pairs). To mitigate sequence redundancy, we clustered protein sequences at a sequence identity threshold of 80% using MMSeqs2 [48] (threshold chosen to optimize out-of-sample performance, see SI Appendix and Fig. S4), reducing the dataset to ∼7 million sequences and ∼9 million reaction-enzyme pairs. We then trained a model to be used for experimental inference, which we call *Horizyn-1* (see Methods). Analysis of *Horizyn-1*’s score distribution revealed a strong divergence between scores for positive and negative reaction-enzyme pairs in the training set, allowing us to use thresholds based on these score distributions to identify high-confidence predictions (see Methods, Fig. S5). This model was applied to predict activity for three experimental use cases: (i) identifying enzymes for orphan reactions, (ii) predicting promiscuous enzyme activity, and (iii) discovering enzymes for non-natural reactions.

### Experimentally de-orphaning reactions and enzymes with *Horizyn-1*

As a first experimental test case for *Horizyn-1*, we sought to identify enzymes for orphaned biochemical reactions in the Rhea database. Orphaned reactions are those that have been experimentally observed or inferred to exist, but for which a catalyzing protein has not yet been annotated [47], thus providing a strong test case for enzyme discovery. We initially focused on the 6-aminohexanoate aminotransferase activity (RHEA:58200, EC 2.6.1.116) observed in *Arthrobacter* sp. KI72, a nylon-degrading bacterium (Fig. 2A) [49]. We queried *Horizyn-1* with the 6-aminohexanoate aminotransferase reaction against all protein coding sequences (*n* = 4,173) from this genome (NCBI accession GCF 002049485.1) (Fig. 2B, Fig. S6). We identified two sequences (WP 021472307.1 and WP 079941468.1) as high confidence candidates. These proteins were previously annotated as a 4-aminobutyrate–2-oxoglutarate transaminase (GabT, EC 2.6.1.19) and an aminotransferase class III-fold pyridoxal phosphate-dependent enzyme (EC 2.6.1.-), respectively, via the NCBI prokaryotic genome annotation pipeline [50] (Fig. 2B). Subsequent expression, purification, and biochemical assay of these two proteins confirmed significant 6-aminohexanoate aminotransferase activity relative to the uncatalyzed reaction (Fig. 2C).

**FIG. 2.**
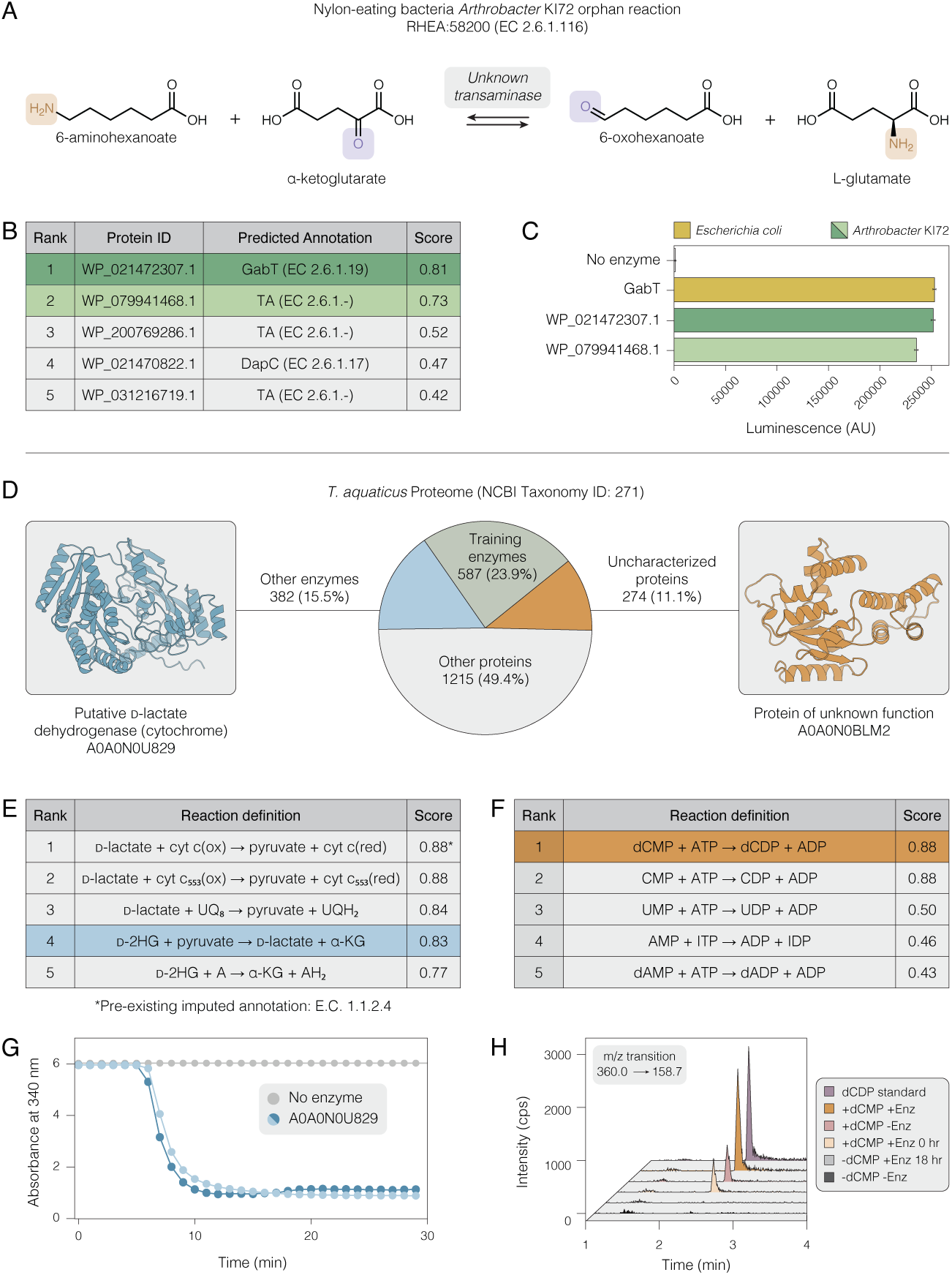
De-orphaning reactions and identification of promiscuous activity in uncharacterized enzymes. **(A)** The orphan 6-aminohexanoate aminotransferase reaction (RHEA:58200) from the nylon degradation pathway in *Arthrobacter* sp. KI72, in which 6-aminohexanoate undergoes transamination with *α*-ketoglutarate as the keto acid acceptor. **(B)** Top 5 candidate proteins from the *Arthrobacter* sp. KI72 proteome predicted by *Horizyn-1* to catalyze the reaction in (A), with putative functional annotations from the NCBI prokaryotic genome annotation pipeline (32). The top two hits, WP 021472307.1 and WP 079941468.1, were selected for validation. **(C)** Experimental validation of 6-aminohexanoate aminotransferase activity. Luminescence, which is proportional to l-glutamate production, is shown for the two candidate *Arthrobacter* enzymes and a known *ω*-transaminase, *E. coli* GabT. **(D)** Pie chart showing the distribution of enzymes, uncharacterized proteins, and other proteins in the *Thermus aquaticus* proteome (NCBI Taxonomy ID: 271). Proteins A0A0N0U829, a putative d-lactate dehydrogenase (cytochrome), and A0A0N0BLM2, a protein of unknown function, were selected for further screening. **(E)** Top 5 Rhea reactions predicted by *Horizyn-1* for the enzyme A0A0N0U829. The model confirms its annotated d-lactate dehydrogenase activity (rank 1) and predicts promiscuous activity for alternative hydride transfer cofactors including *(R)*-2-hydroxyglutarate–pyruvate transhydrogenase activity (d-2HG:*α*-KG) (rank 4). **(F)** Top 5 Rhea reactions predicted by *Horizyn-1* for the uncharacterized protein A0A0N0BLM2, suggesting dCMP and CMP kinase activity. **(G)** Experimental validation of the predicted promiscuous activity for A0A0N0U829. Pyruvate production (for the reverse reaction) is measured via a coupled lactate dehydrogenase assay. NADH concentration (absorbance at 340 nm) is depleted over time only in the presence of the enzyme (replicates, blue lines), confirming transhydrogenase activity. **(H)** Experimental validation of A0A0N0BLM2 activity using LC-MS multiple reaction monitoring (MRM). A peak corresponding to the dCDP product is observed only when dCMP is incubated with the enzyme for 18 h (+dCMP +Enz 18 h), confirming dCMP kinase activity.

The two enzymes identified in *Arthrobacter* sp. KI72 shared significant sequence similarity (43.3% and 43.5%, respectively) with GabT from *E. coli* (P22256) (Fig. 2B). *E. coli* GabT is known to catalyze *ω*-transamination reactions with various substrates, including 4-aminobutyrate and 4-aminopentanoate. Notably, *Horizyn-1* also ranked GabT as the top hit for 6-aminohexanoate aminotransferase in our training set (Fig. S7), suggesting a potential for promiscuous activity. Based on these observations, we purified and tested *E. coli* GabT for 6-aminohexanoate aminotransferase activity, observing higher activity for this reaction than the two *Arthrobacter* sp. KI72 enzymes, and comparable activity to transamination with its annotated substrate, 4-aminobutyrate (RHEA:23352, EC 2.6.1.19) (Fig. 2C, Fig. S7). Furthermore, *Horizyn-1* predicted strong scores for *E. coli* GabT with two additional orphan *ω*-transaminase reactions with alternative substrates, *β*-alanine (RHEA:30699, EC 2.6.1.120) and putrescine (RHEA:23816, EC 2.6.1.82). Experimental testing of GabT with these substrates (along with several others) revealed weak promiscuous activity (Fig. S7), suggesting that *Horizyn-1* may be capable of predicting promiscuous activities for well-characterized enzymes.

We next sought to explore whether *Horizyn-1* could be used to predict the promiscuity of less well-characterized enzymes. To this end, we identified protein sequences from the model thermophilic bacterium *Thermus aquaticus* (NCBI Taxonomy ID: 271) that were identified as enzymes (i.e., had an assigned EC number), but were not included in our training data (Fig. 2D). We searched for sequences that had no more than 50% sequence similarity to any enzymes in our training dataset (*n* = 382), and queried *Horizyn-1* for each sequence against all Rhea reactions (*n* = 15,814). After filtering predictions for reactions that corresponded to previously annotated EC classes, we identified 818 high-scoring (*>*0.80) hits for promiscuous or alternative activity. Upon inspection, we found several high-scoring reactions for a hypothetical d-lactate dehydrogenase (A0A0N0U829, EC 1.1.2.4) (Fig. 2E). The prediction of *Horizyn-1* for the top-scoring reaction for this enzyme (RHEA:13521) was consistent with the result from homology-based models. However, we found that this enzyme also scored highly for *(R)*-2-hydroxyglutarate–pyruvate transhydrogenase (RHEA:51608, EC 1.1.99.40) activity, suggesting this enzyme was promiscuous for alternative electron/donor pairs when oxidizing *(R)*-lactate (Fig. S8). To test for transhydrogenase activity for this enzyme, we expressed, purified, and developed a coupled assay, measuring pyruvate production via a coupled lactate dehydrogenase assay (see Methods). We observed significant depletion of NADH in the presence of this enzyme, suggesting that A0A0N0U829 can act as a *(R)*-2-hydroxyglutarate–pyruvate transhydrogenase (Fig. 2G).

As a more challenging test case, we sought to test whether *Horizyn-1* could be used to predict the activity of completely uncharacterized enzymes. To this end, we identified protein sequences from *T. aquaticus* that were both labeled as an “uncharacterized protein” and contained no homologous sequences in our training data (*n* = 274, see Methods). We then queried *Horizyn-1* for these sequences against all Rhea reactions (*n* = 15,814), and found 61 high scoring (*>*0.80) pairs of reactions and proteins. Upon inspection, we found that the uncharacterized protein A0A0N0BLM2 scored highly for cytidine monophosphate (CMP) and deoxycytidine monophosphate (dCMP) kinase activities (RHEA:11600, RHEA:25094) (Fig. 2F, Fig. S8). We recombinantly expressed and purified this protein, and measured (d)CDP production using LC-MS multiple reaction monitoring (MRM) (Methods). We were able to confirm production of dCDP, but not CDP, suggesting that *Horizyn-1* may experience difficulty discriminating between very similar substrates (Fig. 2H, Fig. S9).

We attempted to replicate *Horizyn-1*’s function predictions for both *T. aquaticus* enzymes using traditional homology methods [6, 48], structural comparisons [52], and modern deep learning tools [36, 55]. While some methods were able to predict the *(R)*-2-hydroxyglutarate–pyruvate transhydrogenase activity of A0A0N0U829, they did so with significantly lower confidence than *Horizyn-1*. However, none of the other methods identified the dCMP kinase activity of A0A0N0BLM2 (see SI Appendix, Table S7). Interestingly, the most similar cytidylate kinase in our training data shared only 28% sequence similarity to A0A0N0BLM2, suggesting that *Horizyn-1* may be useful for predicting promiscuous activities even for sequences with low homology to previously characterized enzymes.

### Enzyme discovery for non-natural biochemical reactions

The experimental confirmation above has been applied to cases where we have screened large libraries of reactions and searched for enzyme-reaction pairs with high model scores. To assess the utility of *Horizyn-1* for an enzyme discovery campaign, we chose to query reactions recently reported to be of interest in the synthesis of non-canonical amino acids (ncAAs) used in pharmaceutical manufacturing [12]. These reactions use so-called “smart” amine donors, which undergo cyclization after amine transfer to the corresponding keto acid acceptor. This cyclization provides a greater thermodynamic driving force than native amine donors such as glutamate (Fig. 3A). Although prior literature reported the feasibility of such reaction types, the specific transaminases capable of performing these reactions were not reported [12]. We used *Horizyn-1* to screen all enzymes in our screening set to generate a panel for experimental validation.

**FIG. 3.**
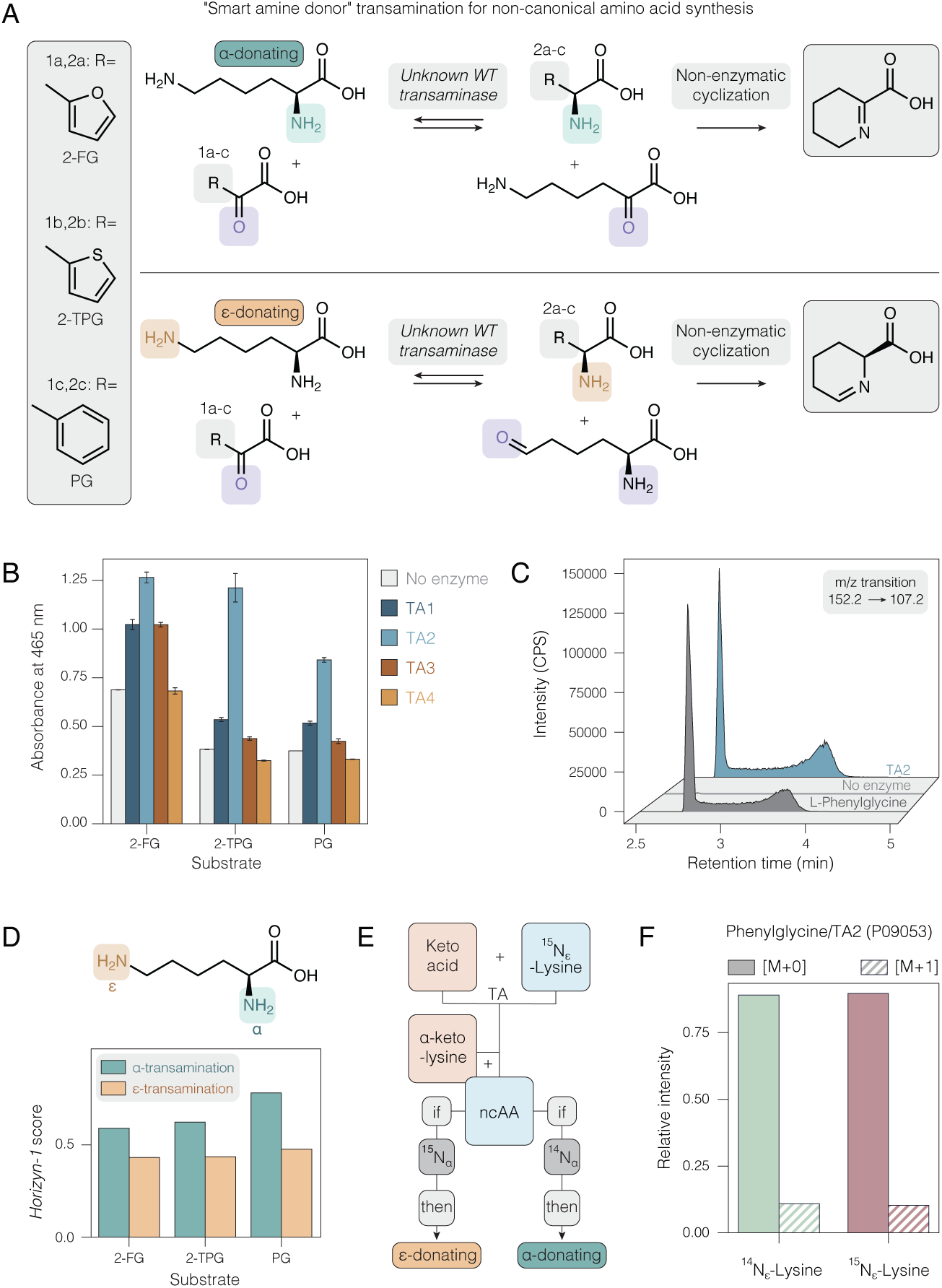
Discovery and mechanistic validation of transaminases for non-canonical amino acid synthesis. **(A)** Schematic of “smart amine donor” transamination for synthesizing non-canonical amino acids (ncAAs). l-lysine can act as an amine donor from either the *α*- or *ε*-position. The resulting keto-acid byproduct undergoes non-enzymatic cyclization, providing a thermodynamic driving force for the reaction. **(B)** In vitro activity screening of four candidate transaminases (TA1: Q9HUI9, TA2: P09053, TA3: Q9I6J2, TA4: O50131) prioritized by *Horizyn-1*. Activity was measured using a colorimetric assay with three different keto acid substrates: 2-furylglyoxylic acid (2-FG), 2-thiopheneglyoxylic acid (2-TPG), and phenylglyoxylic acid (PG). The enzyme TA2 (P09053) demonstrated the highest activity. **(C)** Confirmation of l-phenylglycine production by TA2 via multiple reaction monitoring (MRM). The extracted ion chromatogram for the reaction product (blue trace, *m/z* = 107.2) matches the retention time of an authentic l-phenylglycine standard (gray trace). CPS = counts per second. **(D)** *Horizyn-1* scores for *α*- vs. *ε*-amino-driven transamination for TA2. For all three keto acid substrates, the model predicts that the *α*- donation pathway is strongly favored. **(E)** Experimental design to distinguish between the two potential mechanisms using lysine labeled with a heavy isotope at the *ε*-amine (^15^N*ε*-labeled l-lysine). Label incorporation into the final ncAA product would indicate that the reaction proceeds via *ε*-transamination. **(F)** Relative intensity of unlabeled (*M* +0) and labeled (*M* +1) l-phenylglycine produced by TA2 using either standard ^14^N*ε*-labeled l-lysine or ^15^N*ε*-labeled l-lysine. The relative intensities are computed from summed ion intensities from multiple reaction monitoring (MRM) fragments. The result, showing negligible label incorporation from ^15^N*ε*-labeled l-lysine, confirms that transamination occurs predominantly from the unlabeled *α*-amine, validating the model’s prediction in (D).

We chose three non-canonical keto acids—phenylglyoxylic acid, 2-thiopheneglyoxylic acid, and 2-furylglyoxylic acid—which can be used to synthesize phenylglycine, 2-thienylglycine, and 2-furylglycine, respectively. Two pathways are possible wherein the amine in either the *α*- or *ε*-position can act as the transamination donor (Fig. 3A). We queried *Horizyn-1* for all six reaction variants against all ∼7 million enzymes in our screening set. We then constructed a panel of six transaminases with high scores for all three substrates and at least one of the *α*- or *ε*-donating reaction variants.

Of the six enzymes in our panel, we were able to express and purify four. We screened these four enzymes for activity on all substrates using a colorimetric assay, in which cyclized lysine-derived products react with *o*-aminobenzaldehyde to form compounds with absorbance between 450 and 465 nm (Fig. S10, Methods). We observed significant activity for one enzyme, TA2 (P09053), and minor activity for two others on all three substrates (Fig. 3B). We confirmed the production of the expected non-canonical amino acids via LC-MS MRM and purchased standards; for example, we observed a characteristic transition from 152.2 *m/z* to 107.2 *m/z* for PG (Fig. 3C) as well as characteristic transitions for the other ncAAs (Fig. S11) [17, 23, 25].

Finally, we tested the ability of *Horizyn-1* to distinguish between the *α*- or *ε*-donating reaction variants. For TA2, scores were higher for *α*-transamination compared to *ε*-transamination for all three substrates, suggesting that the mechanism of TA2 transamination is predominantly from the *α*-position (Fig. 3D). We devised an isotope-labeling study to confirm this prediction by supplying ^15^N*_ε_*-labeled l-lysine and measuring the relative intensities of the monoisotopic (*M* + 0) versus heavy isotope (*M* + 1) of l-phenylglycine (Fig. 3E). Consistent with model predictions, we observed comparable heavy versus monoisotopic intensities when supplying either ^15^N*_ε_*-labeled l-lysine or ^14^N*_ε_*-labeled l-lysine, confirming that transamination predominantly stems from the *α*-amine.

### Efficient inclusion of new experimental data rapidly yields predictive models for new reaction classes

Enzyme discovery often requires venturing into new and unexplored areas of chemistry. To assess the model’s ability to generalize to such cases, we tested the model on more challenging splits of our training and evaluation data designed to decrease the attributes shared between the two sets (see SI Appendix). As expected, performance degraded when reactions were stratified to ensure low maximum similarity between training and test reaction sets via fingerprint similarity thresholds or reaction templates (Fig. S12). To more directly assess this relationship between model performance and reaction novelty, we used our best model trained on the training split of the development dataset (Fig. 1B). For each reaction in the development test split, we then correlated the rank of the top-scoring positive enzyme with its distance to the closest reaction in the training split. We found a modest correlation for distances computed using chemical fingerprints (*r* = 0.29), but a substantially stronger one when using the distance within the model’s shared embedding space (*r* = 0.60) (Fig. S13).

This degradation in performance for out-of-sample reactions underscores the critical need to incorporate data from diverse reaction classes to achieve robust generalizability. However, the quantity of experimental data from a new class necessary for accurate prediction remains unclear. To investigate this, we fine-tuned *Horizyn-1* using data from a novel reaction class, ene-reductases (EREDs) (Fig. 4A), and evaluated the model’s resulting performance specifically for that class.

**FIG. 4.**
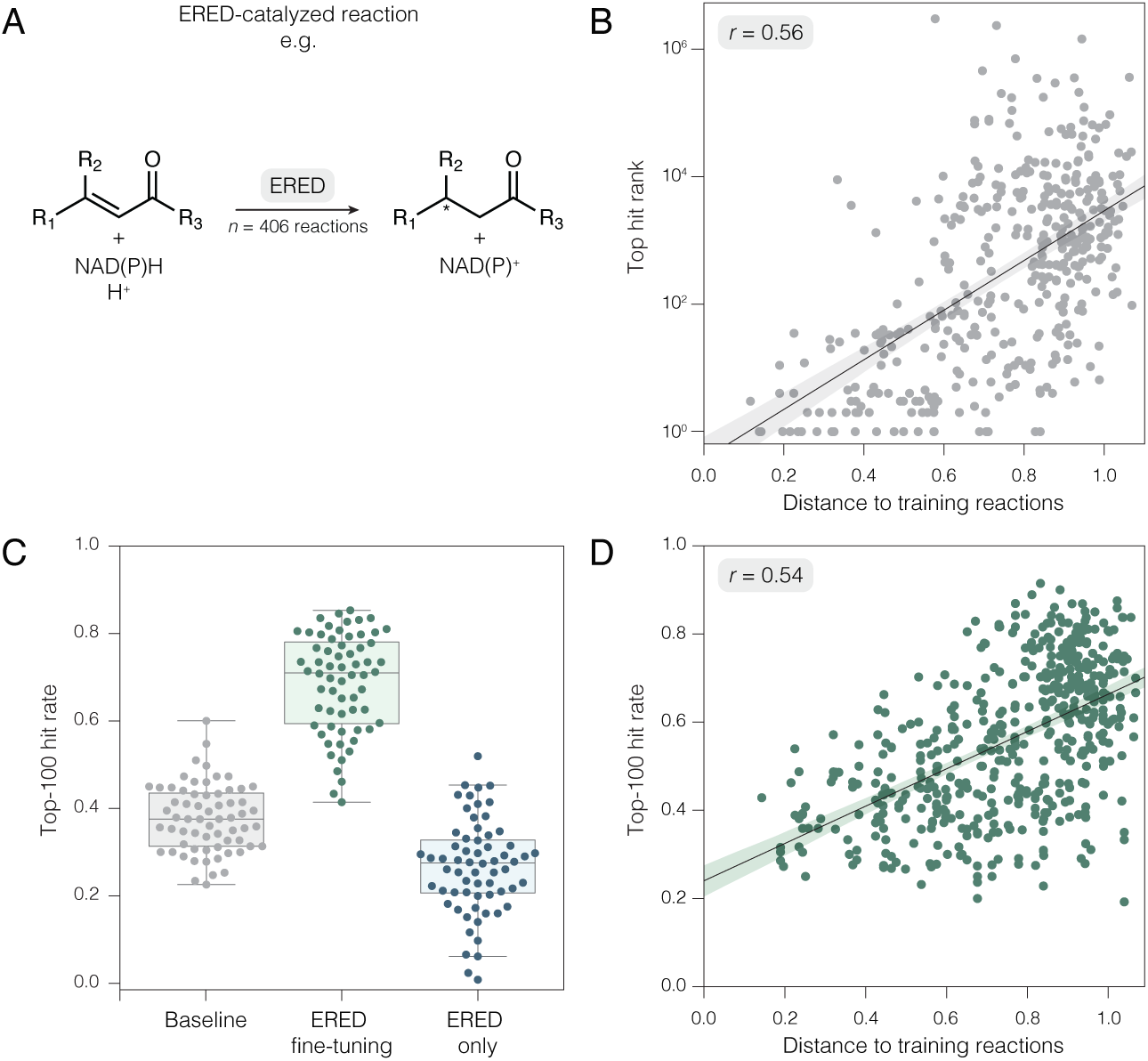
Fine-tuning boosts performance on novel reaction classes. **(A)** Representative ERED-catalyzed reaction involving NAD(P)H-dependent reduction of a C=C bond. This reaction class was not present in the original training data and was used to simulate discovery in a new area of chemistry. **(B)** Baseline model performance on the novel ERED reactions. The rank of the top positive hit (*y*-axis) for each reaction is plotted against its distance from the training set (*x*-axis), measured via the minimum distance to any reaction in the training set within the model’s shared embedding space. Model performance degrades as reactions become more novel. **(C)** Fine-tuning on just 10 new ERED reactions provides a substantial performance boost, increasing the median top-100 hit rate from 38% (baseline) to 71%. Both of these results are a significant improvement over a model trained only on ERED data, which had a median hit rate of 28%. **(D)** Fine-tuning performance (top-100 hit rate) after training on a single ERED reaction plotted against its distance to the training set. The strong correlation demonstrates that fine-tuning on more novel reactions yields greater performance improvements, suggesting an active learning approach to prioritize data collection. In panels (B) and (D), *r* corresponds to the Pearson correlation coefficient with 95% confidence intervals represented as shaded regions.

For this study, we manually curated 406 novel ERED reactions and 903 enzyme-reaction pairs from the literature (see Methods). Consistent with prior results, we found that the baseline inference model’s performance on reactions from this new class strongly correlated with distance from the training set (Fig. 4B, Fig. S12). Next, we fine-tuned the *Horizyn-1* inference model using small, random subsets of these ERED reactions and assessed performance on corresponding held-out validation sets (see Methods). We found that fine-tuning with as few as 10 new ERED reactions substantially improved performance; the median top-100 hit rate increased from 38% to 71% after fine-tuning (Fig. 4C).

To explore the lower limits of data requirement, we also measured the impact of fine-tuning *Horizyn-1* on single ERED reactions. Remarkably, fine-tuning with just one reaction could, in some cases, markedly increase model performance. For example, using only the reduction of (2-nitro-vinyl)-benzene to (2-nitroethyl)-benzene for fine-tuning resulted in a top-100 hit rate of 84% on the ERED test set. However, this effect was highly variable, with post-fine-tuning hit rates ranging from 26% to 84%. We found that the magnitude of this performance increase correlated with the novelty of the single reaction used for fine-tuning relative to the training set (*r* = 0.54) (Fig. 4D, Fig. S14).

Altogether, these results suggest that acquiring even a small number of enzyme-reaction pairs is an effective strategy for rapidly enhancing predictive skill within a new reaction class. Crucially, *Horizyn-1*’s ability to identify reactions that are most distant from its training data provides a mechanism to proactively design experiments that will be most informative for model improvement. This points toward an efficient, model-guided active learning strategy to systematically expand the model’s predictive domain.

## DISCUSSION

The full extent of enzyme diversity and promiscuity for both natural and non-natural biochemical reactions likely far exceeds that reported in literature. With increasing volumes of genomic data, machine learning models like *Horizyn-1* are essential for exploring this diversity, yet their true utility can only be confirmed through rigorous, prospective experimental validation. Such forward-looking assessment, which moves beyond the retrospective benchmarks common in machine learning, is vital for establishing the strengths and limitations of these tools in guiding discovery.

Our findings demonstrate the utility of the dual-encoder contrastive approach employed by *Horizyn-1* for critical biocatalysis tasks, including de-orphaning reactions (Fig. 2A–C), characterizing enzyme promiscuity (Fig. 2D–F), and enabling broader biocatalyst discovery for non-natural reactions (Fig. 4). Crucially, neither standard homology-based approaches like MMSeqs2 or BLAST nor machine learning-based approaches like CLIPZyme were able to replicate the predictions we experimentally verified in this paper for any of these tasks (Table S6, Table S7, Table S8).

Our model is designed to maximize predictive power based on limited available training data. While we sought to maximize our training dataset size, it remains modest compared to those in other domains [40]. By leveraging ProtT5, a 3-billion-parameter protein language model, we harness the knowledge learned from the 10^9^ protein sequences in its pre-training corpus [14]. However, no model of comparable scale exists for chemical reactions, and existing pre-trained models like rxnfp lack sufficient coverage of biochemical transformations [46]. Consequently, we found that a simple fingerprint-based approach exhibits superior performance compared to a reaction model trained from scratch (Fig. 1, Table S5). A likely explanation is that deep learning models struggle to learn informative features from a dataset that is both small and chemically diverse.

Although the model performs well, its capabilities are currently constrained by data availability, with opportunities for improvement through data scaling (Fig. 1E) and fine-tuning (Fig. 4). Notably, model performance degrades for reactions that are increasingly dissimilar to the training data (Fig. 4B, Fig. S12, Fig. S13), while training on even small amounts of novel, sufficiently out-of-sample data can significantly enhance performance (Fig. 4C–D). This result underscores the importance of prioritizing reaction diversity over sheer data volume when collecting new data to improve model performance. Furthermore, distances within *Horizyn-1*’s shared embedding effectively quantify biochemical novelty (Fig. 4B,D, Fig. S13, Fig. S14). This capability could be leveraged to guide the generation of maximally informative data, aiming to build a broadly proficient model [8]. This points toward a lab-in-the-loop active learning strategy that could guide enzyme discovery, dynamically balancing the exploitation of high-confidence predictions with the exploration of novel targets to efficiently expand the model’s knowledge of chemical space.

Beyond data availability, several other limitations constrain predictive accuracy. For instance, the model associates entire enzyme sequences with reactions, without distinguishing the specific domains that catalyze reaction steps or accounting for regulatory elements. Furthermore, current chemical representations are a challenge; reaction SMILES are insufficient for complex molecular contexts like protein-based metabolites (e.g., ferredoxins) or oligonucleotides.

Finally, *Horizyn-1* is primarily designed to retrieve catalytically plausible wild-type enzymes (activity enrichment), not to predict quantitative kinetic parameters or mutant activity, as it lacks access to explicitly labeled negative examples or kinetic measurements (e.g., *k*_cat_, *K*_m_). Nevertheless, performance on enzyme promiscuity benchmarks is encouraging (see Fig. S15): the model discriminates between reactive and non-reactive substrates with an average precision of 28% (compared to an 18% random baseline) and predicted scores exhibit positive correlations with catalytic activity (Spearman *ρ* ≈ 0.16–0.5 for three diverse catalysis datasets). These results suggest that the model captures meaningful functional signals beyond simple retrieval, highlighting clear opportunities for future development as larger and more diverse datasets become available.

Despite recent advances in generative enzyme design [29, 53], enzyme discovery combined with directed evolution remains the gold standard for generating enzymes with novel functionality [2]. *Horizyn-1* promises to accelerate this established discovery pipeline and potentially serve as a foundational model for generative enzyme design tasks, effectively bridging the gap between discovery and *de novo* creation.

## METHODS

### Datasets and Preparation

#### Development Dataset

Our dataset used for model development was curated by combining Rhea (release 131) [3], a collection of expert-curated biological reactions linked to enzymes, and Swiss-Prot (release 2023 05) [51], the manually reviewed, high-quality subset of UniProt. We selected reaction-enzyme pairs from Rhea where the associated enzyme sequence was present in Swiss-Prot. We further excluded any Rhea reactions lacking valid SMILES representations for all participants. The final development dataset contains 11,932 unique reactions, 216,132 unique protein sequences, and 294,166 reaction-enzyme pairs.

#### Inference Dataset

To construct the dataset for training our final inference model, we expanded upon the development dataset by incorporating two additional sources, yielding a final dataset derived exclusively from open-source repositories. First, we added EnzymeMap v2 [21, 22], which contributed 19,140 reactions, 15,308 protein sequences, and 55,163 reaction-enzyme pairs primarily curated from BRENDA [7]. In contrast to Rhea, EnzymeMap contains many reactions that are not naturally occurring.

Second, we augmented the protein sequences using the TrEMBL subset of UniProt (release 2023 05) [51], adding 26,883,845 computer-annotated sequences. While generally less reliable than Swiss-Prot, TrEMBL introduces significant sequence variation and provides additional indirect links between Rhea reactions lacking overlapping Swiss-Prot enzymes. The resulting dataset comprised 31,101 unique reactions, 27,099,893 protein sequences, and 34,385,290 reaction-enzyme pairs.

Finally, we clustered the combined set of all protein sequences using MMseqs2 with a minimum sequence identity threshold of 80% (--min-seq-id 0.8) [48] (https://github.com/soedinglab/MMseqs2). This threshold was chosen based on the performance of the development model on the test set after clustering sequences in the training set (see SI Appendix and Fig. S4). This step substantially reduced sequence redundancy and created further indirect reaction links via enzymes with highly similar sequences. The resulting inference dataset was reduced to 7,063,237 clustered protein sequences and 8,897,870 reaction-enzyme pairs.

#### Arthrobacter Genome Annotation

Protein coding sequences from *Arthrobacter* sp. KI72 (*Paenarthrobacter ureafaciens* KI72), a nylon-eating bacterium, were obtained from the NCBI FTP server using the accession GCF 002049485.1, with annotations derived from the NCBI Prokaryotic Genome Annotation Pipeline [50]. Protein coding sequences (and corresponding EC annotations) for *Thermus aquaticus* were obtained from the UniProt reference proteome for Taxonomy ID: 271. For enzyme promiscuity analysis, we screened all enzymes with no annotation in Rhea (release 131), and had at most 50% sequence similarity to enzymes in our training data. Sequence similarity was computed using MMseqs2 [48] using default settings.

#### Ene-Reductase (ERED) Dataset

The ene-reductase dataset was compiled from reactions available in the BRENDA database (EC classes 1.3.1.-, 1.6.99.-) [7] and the public version of the RetroBioCat ERED dataset [16]. This collection was supplemented with additional manual curation from primary literature sources. This dataset totaled 543 reactions, 299 proteins, and 1,136 pairs. To ensure reactions were distinct from the inference dataset, they were filtered to remove those with identical fingerprints (see Structural Fingerprint Clustering Protocol). This process resulted in a final dataset of 406 reactions, 221 proteins, and 903 pairs.

### Reaction SMILES Standardization and Augmentation

To ensure representation consistency across data sources and compatibility with the reaction fingerprinting algorithms we investigated, we used RDKit [28] to standardize all reaction SMILES strings in the development, inference, and ene-reductase datasets with the following steps:

1. Convert hypervalent N, P, Cl, Br, I to zwitterionic structures: `rdmolops.SanitizeMol` with the `SANITIZE CLEANUP` flag
2. Neutralize acidic and basic sites by adjusting ionizable protons: `rdMolStandardize.Uncharger`
3. Detach metals by converting covalent metal-heteroatom bonds to ionic bonds: `rdMolStandardize.MetalDisconnector`

We found these steps to be sufficient for reactions sourced from Rhea and EnzymeMap. However, many of the ene-reductase-catalyzed reactions were manually curated directly from literature, resulting in many cases where the same reaction was represented with different choices of SMILES. Therefore, we applied two extra standardization steps for this dataset:

4. Generate canonical tautomer: `rdMolStandardize.TautomerParentInPlace`
5. Kekulize aromatic structures: `rdmolops.Kekulize`

Because many of the reaction representations we investigated were not invariant to reaction direction, we augmented reactions by including separate copies of each reaction proceeding in both the forward and backward directions. This resulted in each reaction-enzyme pair being duplicated with both directions during training and for computing performance metrics. We always included both copies of each reaction in the same split during model training and evaluation to avoid contamination.

#### Structural Fingerprint Clustering Protocol

Our default clustering protocol quantifies similarities between reactions by computing Tanimoto coefficients between their structural fingerprints. First, we applied our standardization protocol to all reaction SMILES (see Reaction SMILES Standardization and Augmentation). Next, we used the RDKit library [28] to generate 2048-bit extended-connectivity fingerprints with radius 3 (ECFP6) for each molecule via the rdFingerprintGenerator.GetMorganGenerator class. We enabled options to incorporate chirality, bond orders, and ring membership information. For each reaction, we consolidated the reactant and product fingerprints into separate binary vectors using a bitwise OR operation and then concatenated these vectors to create a single 4096-bit reaction fingerprint. The concatenation order reflects the reaction direction, with the reactant fingerprint preceding the product fingerprint. To ensure directional invariance, we computed Tanimoto coefficients between two reactions for both the forward and reverse directions of each, taking the maximum value from the four possible comparisons. We grouped reactions with a Tanimoto coefficient of 1.0 into the same cluster. Finally, we randomly assigned these clusters to the training, validation, or test sets based on the desired split ratio.

### Model Architecture and Training Protocol

#### Reaction Representations

We evaluated three primary categories of chemical reaction encodings: (i) algorithmically generated graph-based representations, (ii) machine-learned graph-based representations, and (iii) text-based representations. Because text-based approaches typically utilize SMILES strings, they are often sensitive to canonicalization artifacts, as a single compound may correspond to multiple valid SMILES strings. In contrast, graph-based approaches circumvent this ambiguity by deriving features directly from the invariant molecular graph structure.

##### Extended-connectivity Fingerprints (ECFP6)

To capture molecular structure, we generated extended-connectivity level 3 fingerprints (ECFP6) [42, 45] as specified in the Structural Fingerprint Clustering Protocol. For use in *Horizyn-1*, we used fingerprints composed of 1024 bits, with 512 bits allocated for each of the reactant and product subcomponents.

##### DRFP Fingerprints

To incorporate the approximate reaction mechanism and bond changes, we employed a different fingerprint known as DRFP (Difference Reaction Fingerprint) [38]. We used the default DRFP parameters (min radius=0, radius=3, rings=True, root central atom=True, include hydrogens=False), with the exception of setting the output fingerprint dimension to 1024 bits.

##### ECFP6/DRFP Composite Fingerprints

We constructed composite reaction fingerprints by concatenating the ECFP6 and DRFP fingerprints described above. This allowed us to capture both molecular structure and reaction transformations. The final reaction fingerprint used in our study is the direct concatenation of the 1024-bit ECFP6 structure fingerprint and the 1024-bit DRFP transformation fingerprint, yielding a total size of 2048.

##### Chemprop

Chemprop [20] is a graph neural network model based on the Directed Message Passing Neural Network (D-MPNN) architecture, originally developed for predicting molecular properties. While often applied to single molecules, Chemprop’s framework can be adapted for reaction modeling, particularly as it can handle multi-component systems.

In our implementation, we used a Chemprop-style architecture consisting of three layers of directed message passing. To obtain a fixed-size representation for each reaction system (reactants and products), we aggregated the node features learned by the message-passing layers using a global pooling operation (e.g., summing node features) followed by an MLP projection. This resulted in a single vector representation in the shared embedding space. This Chemprop model was trained from scratch using our reaction dataset as part of the end-to-end training process for the contrastive model.

##### rxnfp

The rxnfp model [46] is a pre-trained BERT-style transformer encoder. This model was trained on 2.6 million reactions from the Pistachio dataset using a Masked Language Modeling (MLM) objective, where it learned to predict masked tokens within reaction SMILES strings.

For our dual contrastive model, we used the pre-trained rxnfp model to process reaction SMILES strings. Following the fine-tuning methodology proposed by the original authors, we extracted embeddings from the final classification token ([CLS]) of the transformer output. This process yields a single vector representation for each reaction, which serves as the input embedding for the subsequent multilayer perceptron (MLP) head in our model. We investigated two strategies: using the frozen pre-trained weights and fine-tuning (unfreezing) the rxnfp weights during the training of our contrastive model.

#### Enzyme Representations

To generate enzyme vector embeddings, we employed two pre-trained protein language models (pLMs): ProtT5 and ESM-2. Inspired by transformer models in the natural language processing domain, pLMs treat protein sequences analogous to text with amino acids acting as the vocabulary. Both models were trained on a Masked Language Modeling (MLM) objective, where the goal is to predict the identity of masked amino acid tokens from their context. This self-supervised training objective, when applied to a diverse collection of sequences, enables the models to learn rich embeddings of both structure and function.

##### ProtT5

ProtT5 is a 3-billion-parameter encoder-decoder transformer pre-trained on large sequence datasets totaling over 2.1 billion sequences [14]. To create a fixed-size (1024-dimensional) embedding for each protein, we extracted the final decoder output embeddings for each token (residue) and applied mean pooling. Sequences longer than the model’s limit (5,000 residues) were processed in chunks, and their embeddings were averaged. We precomputed the embeddings before training for efficiency.

##### ESM-2

The ESM-2 family of models employs encoder-only (BERT-style) transformers pre-trained on over 65 million sequences [30]. While we explored multiple configurations, including alternative embedding extraction strategies and end-to-end fine-tuning (see SI Appendix), our primary approach used static, pre-computed embeddings. These were generated from the final hidden state of the [CLS] token, which aggregates sequence-level information. We generated these embeddings for both the 650M and 3B parameter versions of ESM-2; sequences exceeding the 1,500-residue input limit were truncated.

#### Dual Encoder Architecture

Our model maps reaction and enzyme representations into a shared embedding space via two multilayer perceptrons (MLPs) that act in parallel. Each MLP accepts representation-specific input dimensions and uses two 4096-dimensional hidden layers with ReLU activation. The resulting embedding vectors are L2-normalized, and cosine similarity (dot product) is used to compute their distances.

#### Contrastive Learning and Negative Sampling

In contrastive learning a model is trained to embed pairs of objects in a shared embedding space, maximizing the similarity of associated (positive) pairs while minimizing that of unassociated (negative) pairs. A critical component of this framework is the strategy for negative sampling. While some approaches utilize hard negative mining to specifically target negative pairs that are difficult for the model to distinguish [33], we employ a Noise Contrastive Estimation (NCE) framework [32]. In this context, the model is trained to distinguish positive pairs from a background distribution of noise, which takes the form of random unannotated reaction-enzyme combinations constructed from the dataset.

To implement this efficiently, we utilize in-batch negative sampling as demonstrated in CLIP [40], where all non-paired elements within a mini-batch serve as negatives for one another. This strategy avoids the computational overhead of mining specific hard negatives while facilitating incredibly deep sampling—exposing the model to a vast number of negative examples over the course of training. This high volume of negatives is thought to be beneficial for learning robust representations [9].

#### Maximum Likelihood Noise Contrastive Estimation (MLNCE) Loss

In this work, we adapt the objective function utilized in CLIP [40] to formulate a rigorous Maximum Likelihood Estimation (MLE). By grounding the contrastive loss in a probabilistic framework, we utilize the Maximum Likelihood Noise Contrastive Estimation (MLNCE) loss. The complete derivation of this objective is provided in the SI Appendix.

#### Model Evaluation

Model performance was assessed using an evaluation set containing reactions, associated enzymes, and a broader screening set. For each reaction, the primary evaluation measured how effectively the model ranked known catalyzing enzymes (true positives) higher than other enzymes within a screening set. For a particular reaction, positive enzymes were those derived from reaction-enzyme pairs involving that reaction. Performance was quantified using several standard information retrieval metrics. Each metric was averaged over each reaction, proceeding in both the forwards and backwards directions.

Top-*k* hit rate, which measures whether at least one of the true positives appears within the top *k* ranked results returned by the model, was chosen to reflect the practical, real-world use case of identifying relevant enzymes from a large pool with *k* values of 1, 10, 100, and 1000. *R*-precision and average precision offer more stringent assessments of retrieval quality. *R*-precision is the strictest metric, measuring the fraction of a reaction’s true positive enzymes found within the top *R* ranked results, where *R* equals the total number of true positives for that reaction. Average precision provides a less strict, but more comprehensive measure of how highly positive enzymes are ranked relative to others across the entire list, offering greater sensitivity to the ranking of all relevant items and being less susceptible to class imbalance issues compared to metrics like AUROC. Additionally, Boltzmann-Enhanced Discrimination of Receiver Operating Characteristic (BEDROC) and Enrichment Factor (EF) were included, primarily for comparison purposes with the CLIPZyme results. BEDROC is a metric that heavily weights the ranking of true positives found early in the list, assessing initial retrieval performance. Enrichment Factor (EF) measures the concentration of true positives within a specific top fraction of the ranked list compared to what would be expected from a random ranking. For consistency with the CLIPZyme results, we report BEDROC scores with *α* equal to 20 and 85, and EF scores for the top 5% and 10% of predictions.

#### General Training Protocol

All models were trained using PyTorch v. 2.0.0 [37] and PyTorch Lightning v. 2.3.3 [15]. We employed the AdamW optimizer with parameters (*β*_1_, *β*_2_) = (0.9, 0.999), a weight decay of 10^−2^, and a fixed learning rate of 5 × 10^−5^. A default batch size of 16,384 was used unless otherwise specified. With the exception of the ESM-2 protein language model, all models were trained on a single NVIDIA T4 GPU.

To ensure consistent batch sizes, crucial for the MLNCE loss (see SI Appendix), the final batch of each epoch was padded to the required size. Padding involved randomly duplicating reaction-enzyme pairs from the training data.

We used a fixed inverse temperature parameter *β* = 10 to set the distance scale of our loss function (see SI Appendix). While the original CLIP implementation treats this as a learnable parameter, we observed slightly improved results and more stable training dynamics by fixing its value.

Training duration was determined based on validation set performance, approximately targeting epochs after the point where the top-100 hit rate on a held-out validation set began to plateau, but before the onset of overfitting.

#### Development Model Training and Evaluation

The development model was trained on the development dataset (see Development Dataset), derived from Rhea and Swiss-Prot. The training and test sets were created by splitting on reactions and subsequently filtering the test set to exclude reactions that had fingerprint similarities of 1.0 with any reactions in the training set (see Structural Fingerprint Clustering Protocol). The final training and test splits contained 10,785/1,012 reactions, 192,769/32,100 enzymes, and 257,733/33,996 reaction-enzyme pairs. The model was trained for 100 epochs with batch sizes of 16,384 (default), 256 (Chemprop), and 16 (unfrozen rxnfp). For evaluation, the model identified the correct enzyme for each test reaction from a screening pool of all 216,132 proteins in the development dataset. Results are shown in Fig. 1D.

#### Inference Model Training

The inference model was trained using the inference dataset, derived from Rhea, EnzymeMap, Swiss-Prot, and TrEMBL (see Inference Dataset). For this model, the entire dataset was used as the training set, with all proteins used for the screening set. The model was trained for 200 epochs.

#### Training Set Size Scaling Analysis

To analyze the impact of training set size (Fig. 1E), we first subdivided the training subset of the development dataset (see Development Model Training and Evaluation) into a fixed random 80%/20% training/validation split based on reaction fingerprint clusters (see RStructural Fingerprint Clustering Protocol). To create datasets for each run, the 80% training portion was then systematically subsetted by reaction clusters to achieve the desired training set sizes. All models were trained for 100 epochs and evaluated on the same fixed validation set, using the full development dataset’s protein set as the screening pool.

#### Ene-Reductase Fine-tuning Protocol

The fine-tuning studies (Fig. 4) used a manually curated ene-reductase dataset (see Ene-Reductase Dataset). The data splits for fine-tuning were prepared using the following procedure:

1. Randomly split the ene-reductase dataset into a 90%/10% train/val split based on reaction fingerprint clusters (see Structural Fingerprint Clustering Protocol).
2. Randomly select the desired number of reaction clusters from the training split. Results in Fig. 4C and D used 10 and 1 reaction clusters, respectively.
3. Add an additional 10 random reaction-enzyme pairs from the inference dataset to the training split, with each reaction belonging to a different fingerprint cluster. These additional pairs served to regularize the model during fine-tuning and prevent total collapse of the shared embedding space.

Each training replicate used a different random seed to randomize each of these steps. Model evaluation was performed on reaction-enzyme pairs from the validation split, using a comprehensive screening set that combined all proteins from both the inference and ene-reductase datasets, totaling 7,063,403 proteins. Training for fine-tuning was conducted for 10 epochs. Models were initialized either from scratch with random weights or with the weights of the previously established inference model for fine-tuning. The batch size was set to twice the number of pairs in the final training split; the factor of two accounted for reaction direction augmentation.

### Model Confidence Heuristic

To provide a practical framework for interpreting model outputs in experimental settings, we established a qualitative confidence heuristic based on the *Horizyn-1* score distributions from the inference model training set (Fig. S5C). This approach categorizes predictions into three confidence tiers based on the score distributions of positive and negative enzyme-reaction pairs. The tiers are defined as follows:

- **Low Confidence (Score** *<* **0.10):** Scores falling below the 99th percentile of the negative pair distribution.
- **Moderate Confidence (Score 0.10–0.72):** Scores between the 99th percentile of negative pairs and the 1st percentile of positive pairs.
- **High Confidence (Score** ≥ **0.72):** Scores at or above the 1st percentile of the positive pair distribution. This heuristic is intended to guide the prioritization of candidates for experimental screening.

### Experimental procedures

#### Chemical Reagents

The compounds *α*-ketoglutaric acid sodium salt, lysine, MnSO_4_, Co_3_(PO_4_)_2_, MgCl_2_, ZnCl_2_, *(R)*-lactate, l-lactate dehydrogenase, NADH, UMP, and CMP were purchased from Millipore Sigma. Additional chemicals including 6-aminohexanoic acid, monosodium l-glutamate, ATP, PLP, *o*-aminobenzaldehyde, and 2-thiopheneglyoxylic acid were also sourced commercially (VWR).

#### Cloning and transformation

The native *E. coli gabT* gene (Genbank M88334) encoding the Uniprot P22256 sequence was trimmed to remove the start and stop codons and ordered from Twist Biosciences as a clonal gene inserted between the *Nde*I and *Hind*III restriction sites in pET29b(+). The insertion resulted in a full-length transcript containing an additional seven amino acid linker (KLAAALE), C-terminal His-tag, and stop codon encoded in the plasmid sequence.

DNA sequences for GabT transaminase homologs from *Arthrobacter* sp. KI72 (NCBI sequences WP 021472307.1 and WP 079941468.1), lysine TAs (UniProt: Q9HUI9, P09053, Q9I6J2, O50131), transhydrogenase from *Thermus aquaticus* (UniProt: A0A0N0U829), and dCMP kinase from *Thermus aquaticus* (UniProt: A0A0N0BLM2) were synthesized without the start and original stop codon from Twist Biosciences and modified to contain a C-terminal GGSGSG linker, 6-His tag, and stop codon. Full ORFs were codon optimized for expression in *E. coli*. These sequences were delivered as clonal constructs in pET29b(+) with gene insertion at the *Nde*I and *Xho*I sites.

#### Protein overexpression and purification

Plasmids harboring GabT, GabT transaminase homologs, lysine TAs, and dCMP kinase were transformed into chemically competent BL21(DE3) cells (Agilent Technologies). The plasmid harboring transhydrogenase was transformed into chemically competent OverExpress C41(DE3) cells (Sigma-Aldrich). Cells were plated on LB agar containing 50 *µ*g/mL kanamycin for overnight growth. Single colony transformants were grown aerobically overnight at 37 ^◦^C in 5 mL of LB medium supplemented with 100 *µ*g/mL kanamycin. Cells from the starter cultures were added to 100 mL LB with 100 mg/L kanamycin and cultivated at 37 ^◦^C until OD_600_ reached 0.6–1.0. Expression was induced with 0.2 mM IPTG for 16 h at 25 ^◦^C for GabT. For other enzymes, expression was induced with 0.2 mM IPTG for 16 h at 16 ^◦^C. Cells were then harvested by centrifugation (e.g., 8000 ×*g*, 15 min, 4 ^◦^C) and stored at −20 ^◦^C until use.

Enzymes were purified through Ni-NTA gravity column purification. Cell pellets were thawed and resuspended in one-fifth the volume of the culture with Lysis Buffer (20 mM Tris, pH 7.5, 0.5 M NaCl, 100 *µ*M PLP, 10 mM imidazole, protease inhibitor cocktail (Roche), DNase, and lysozyme) and then lysed by sonication on ice using a microtip on a Branson sonifier at 70% amplitude with five 1 s on / 1 s off cycles, pausing for 30 s on ice between cycles. The resulting lysate was centrifuged for 30 min at 20,000 ×*g*. Clarified lysate was collected and incubated with 500 *µ*L of washed Ni-NTA beads (Thermo Fisher) per manufacturer’s instructions with rocking on a nutator for 30 min.

Lysate was loaded onto a gravity column and the resin was washed with 10 CV of Wash Buffer (20 mM Tris pH 7.4, 0.5 M NaCl, and 20 mM imidazole). The protein was eluted with 10 CV of Elution Buffer (20 mM Tris pH 7.4, 0.5 M NaCl, and 250 mM imidazole). Fractions were collected and analyzed by SDS-PAGE (GenScript). Purified enzymes were dialyzed with 50 mM potassium phosphate pH 8.0 with a 30 kDa MWCO spin column (Cytiva) and stored in 12.5% glycerol at −80 ^◦^C until use. The concentration of purified enzyme was determined by the Pierce BCA assay (Thermo Fisher) per manufacturer’s protocol.

#### Enzymatic reaction and assays

##### Transaminase Assay

*E. coli* GabT and GabT transaminase homologs from *Arthrobacter* sp. KI72 were tested for aminotransferase activity using the amino donor 6-aminohexanoic acid and the amino acceptor 2-oxoglutarate. Reactions were assembled containing 50 mM sodium phosphate, pH 8.0, 1 mM PLP, 100 mM 2-oxoglutarate, and 50 mM of amino donor (6-aminohexanoate, *β*-alanine, or putrescine). To initiate the reaction, purified enzyme was added at a final concentration of 10 *µ*M. For negative control, purified enzyme was replaced with the same volume of 50 mM sodium phosphate buffer. Reactions were performed at room temperature for approximately 16 h. The formation of glutamate in transaminase enzymatic assays was detected by using the Glutamate-Glo™ Assay (Promega) following the manufacturer’s protocol. A luminescence signal indicating glutamate formation was measured by a POLARStar Omega plate reader (BMG Labtech). All data points were measured in duplicate.

##### Lysine Transaminase Assay

Ability to transfer an amino group from lysine to an oxoacid to form a non-canonical amino acid was determined for Q9HUI9, P09053, Q9I6J2, and O50131. Reaction mixtures contained 50 mM sodium phosphate, pH 8.0, 0.4 mM PLP, 5 mM *o*-aminobenzaldehyde, 100 mM ketoacid, and 100 mM l-lysine. Reactions were initiated through addition of 10 *µ*M of purified enzymes and allowed to react at 37 ^◦^C for approximately 18 h. For negative controls, purified enzyme was replaced with the same volume of 50 mM sodium phosphate buffer. Reactions containing lysine as the amino donor were tested colorimetrically, taking advantage of the spontaneous cyclization of products and reactivity with *o*-aminobenzaldehyde to form colored compounds. If the *α*-amine is used as a donor, 6-amino-2-oxohexanoic acid is formed which spontaneously cyclizes to 2-piperideine-2-carboxylate. The piperideine reacts with *o*-aminobenzaldehyde to form a three-membered ring product with an absorbance maximum of 450 nm [17, 23]. Use of the *ε*-amine as a donor results in formation of l-allysine. l-allysine similarly cyclizes to *(S)*-1-piperideine-6-carboxylate and reacts with *o*-aminobenzaldehyde to form a three-membered ring product with an absorbance maximum at 465 nm [23, 25]. Absorbance was measured by a POLARStar Omega plate reader (BMG Labtech). Quantification was performed using an extinction coefficient of 2.8 mM^−1^cm^−1^. One unit of enzyme activity is defined as catalyzing formation of 1 nmol of product per min. All data points were measured in duplicate.

##### Transhydrogenase Assay

Transhydrogenase A0A0N0U829 from *Thermus aquaticus* was tested for transhydrogenase activity. Transhydrogenase reactions contained 50 mM Tris, pH 7.5, 0.1 mM ZnCl_2_ or 1 mM of other divalent cations (MnSO_4_, Co_3_(PO_4_)_2_, MgCl_2_), 50 mM 2-ketoglutarate, 200 mM *(R)*-lactate, and 10 *µ*M purified enzyme. For negative controls, purified enzyme was replaced with the same volume of 50 mM Tris, pH 7.5 buffer. Reactions were run at 55 ^◦^C for approximately 18 h and stopped by heating at 95 ^◦^C for 3 min. Samples were centrifuged for 10 min at 16,000 ×*g* at 4 ^◦^C to remove protein precipitation and the pyruvate formation was determined via l-LDH (Roche) by monitoring loss of absorbance of NADH at 340 nm. The l-LDH assay mixture contained 100 mM Tris-HCl, pH 7.5, 3 mM NADH, 0.1 unit l-LDH, 50 *µ*L reaction mix in a total reaction volume of 200 *µ*L. Absorbance was measured on a POLARStar Omega plate reader (BMG Labtech). All data points were measured in duplicate.

##### Kinase Assay

A0A0N0BLM2 from *Thermus aquaticus* was tested for kinase activity. Reactions contained 50 mM Tris, pH 7.5, 5 mM MgCl_2_, 1 mM ATP, 1 mM CMP or dCMP, 10 *µ*M purified enzyme. For negative controls, purified enzyme was replaced with the same volume of 50 mM Tris, pH 7.5 buffer. Reactions were run at room temperature for ∼18 h. The formation of ADP was detected by ADP-Glo™ Assay (Promega) following the manufacturer’s protocol. CDP and dCDP were detected by LC-MS as described in the LC-MS section. All data points were measured in duplicate.

#### LC-MS

Non-canonical amino acid products of lysine transaminases and dinucleotide products of A0A0N0BLM2 from *Thermus aquaticus* were confirmed via LC-MS consisting of a LC-20AD liquid chromatography system (Shimadzu) and 3200 QTRAP MS/MS (SciEx). Separations were performed on an ACQUITY UPLC HSS T3 column, 100 Å, 1.8 *µ*m, 2.1 × 150 mm (Waters). Mobile phase A was composed of 0.1% formic acid in water, and mobile phase B was acetonitrile with 0.1% formic acid. The gradient was as follows: 0 min, 0% B; 0 to 2.5 min, 4.8% B; 2.5 to 3.5 min, 4.8 to 30% B; 3.5 to 4.10 min, 30 to 100% B; 4.1 to 5.1 min, 100% B; 5.1 to 5.2 min, 100% to 0% B; 5.2 to 10 min, 0% B. The flow rate was 0.2 mL/min. The injection volume was 10 *µ*L and the column was maintained at 40 ^◦^C. Experiments were performed on a triple quadrupole electrospray ionization (ESI) 3200 QTRAP MS/MS (SciEx) operating in positive ion mode. Analytes were fragmented using nitrogen gas as the collision gas and optimized multiple reaction monitoring transitions were recorded for each analyte. Mass spectrometry parameters were as follows: A) l-phenylglycine: ion source gas 1 (GS1) 50.00 arbitrary units, ion source gas 2 (GS2) 50.00 arbitrary units, curtain gas (CUR) 20.00 arbitrary units, ionization spray 5000.00, CAD Medium, declustering potential (DP) 20.00 V, entrance potential (EP) 10.00 V, collision cell entrance potential (CE) 15.30 V, collision cell exit potential (CXP) 4.00 V, source temperature 500 ^◦^C; B) 2-thienylglycine: GS1 50.00, GS2 50.00, CUR 20.00, IS 5000.00, CAD Medium, DP 17.00 V, EP 4.00 V, CE 22.00, CXP 2.50, source temperature 500 ^◦^C; C) 2-furylglycine: GS1 50.00, GS2 50.00, CUR 20.00, IS 5000.00, CAD Medium, DP 20.00 V, EP 10.00 V, CE 30.00, CXP 5.00, source temperature 500 ^◦^C; D) CDP: GS1 50.00, GS2 50.00, CUR 20.00, IS 5000.00, CAD Medium, DP −47.00 V, EP −4.00 V, CE −31.00, CXP −1.00, source temperature 500 ^◦^C; E) dCDP: GS1 50.00, GS2 50.00, CUR 20.00, IS 5000.00, CAD Medium, DP −45.00 V, EP −5.00 V, CE −34.00, CXP −1.36, source temperature 500 ^◦^C. MRM transitions and retention times are listed in Table S11. For phenylglycine, peak areas were integrated between 2.97 and 4.20 min. Note, in Fig. 3, we cannot distinguish between l- and d-enantiomers of the amino acid products because the chromatography was performed on an achiral column, which does not resolve stereoisomers.

## DATA AND SOFTWARE AVAILABILITY

The development dataset is archived on Zenodo (DOI: 10.5281/zenodo.17957034). Code for the Horizyn-1 development model is available on GitHub (https://github.com/dayhofflabs/horizyn).

## ACKNOWLEDGMENTS

We thank Margaux Pinney, Adrian Jinich and Lynn Kamerlin for helpful discussions. We are also grateful to Avi Flamholz for providing valuable comments on the manuscript. We acknowledge support by the Department of Energy Small Business and Innovation Research (SBIR) program through grant number DE-SC0025760. Finally, we thank Rita Clare/Scivetica for assistance with illustrations.

