## Supplementary Information for "Dual-encoder contrastive learning accelerates enzyme discovery"

### Supporting Information for Dual-encoder contrastive learning accelerates enzyme discovery

Jason W. Rocks,[1] Dat P. Truong,[1] Dmitrij Rappoport,[1] Samuel Maddrell-Mander,[1]
Daniel A. Martin-Alarcon,[1] Toni M. Lee,[1] Steven Crossan,[1] and Joshua E. Goldford[1,*]
[1]*Dayhoff Labs, Inc. Cambridge, MA, USA*

#### EC ANNOTATIONS AND CONDITIONING ANALYSIS

##### EC Annotations

EC annotations for the development dataset were curated from the corresponding releases of Rhea and Swiss-Prot. For enzymes, we utilized existing UniProt annotations, covering 98% of enzymes with at least a first-digit EC number and 92% with a full four-digit EC number. For reactions, we first retrieved EC numbers from Rhea, which provided coverage for 47% of the dataset. For the remaining reactions, we attempted to infer EC numbers based on the annotations of all enzymes associated with the reaction in UniProt. This resulted in a total coverage of 91% with at least a first-digit EC number annotation, and 79% with full four-digit classification.

##### EC Conditioning Analysis

For the EC number conditioning analysis (Fig. 1D, Table S3), we ranked proteins for a given query reaction by assigning a score from 0 to 4, corresponding to the number of matching digits in their respective EC numbers. For the EC-only baseline, ties were broken randomly, and final retrieval metrics were averaged over 10 such permutations. To evaluate the retrieval performance when conditioned on EC numbers, we combined the scores using the formula: model score + 2 × EC score. This weighting scheme effectively uses our model to break ties among proteins with identical EC scores.

#### DERIVATION OF MAXIMUM LIKELIHOOD NOISE CONTRASTIVE ESTIMATION (MLNCE) LOSS

To formally derive the training objective, we first establish the notation for the reaction and protein spaces and their representations. Let $\mathcal{R}$ denote the set of reactions, $\mathcal{P}$ the set of enzymes, and $\mathcal{A} \subseteq \mathcal{R} \times \mathcal{P}$ the set of annotated pairs $(r, p)$ where enzyme $p \in \mathcal{P}$ is known to catalyze reaction $r \in \mathcal{R}$. We represent our reaction and enzyme encoders as functions $\vec{\mathbf{f}}_R : \mathcal{R} \to S^{d-1}$ and $\vec{\mathbf{f}}_P : \mathcal{P} \to S^{d-1}$, respectively. These map entities into a shared embedding space defined by the unit hypersphere $S^{d-1} = \{\vec{\mathbf{v}} \in \mathbb{R}^d \mid \|\vec{\mathbf{v}}\|_2 = 1\}$.

We cast reaction-enzyme association as a maximum likelihood estimation problem. Our objective is to maximize the joint probability of drawing annotated reaction-enzyme pairs from the set of all possible pairs. We define the probability of a reaction-enzyme pair $(r, p)$ conditioned on model parameters $\vec{\boldsymbol{\theta}}$ as

$$P(r, p|\vec{\boldsymbol{\theta}}) = \frac{e^{\beta \vec{\mathbf{f}}_R(r;\vec{\boldsymbol{\theta}})\cdot\vec{\mathbf{f}}_P(p;\vec{\boldsymbol{\theta}})}}{\sum_{(r',p')\in\mathcal{R}\times\mathcal{P}} e^{\beta \vec{\mathbf{f}}_R(r';\vec{\boldsymbol{\theta}})\cdot\vec{\mathbf{f}}_P(p';\vec{\boldsymbol{\theta}})}} = \frac{e^{\beta \vec{\mathbf{f}}_R(r;\vec{\boldsymbol{\theta}})\cdot\vec{\mathbf{f}}_P(p;\vec{\boldsymbol{\theta}})}}{Z(\vec{\boldsymbol{\theta}})} \tag{1}$$

where $\beta$ is an inverse temperature parameter scaling the distances in the embedding space. The partition function $Z(\vec{\boldsymbol{\theta}})$ sums over all possible reaction-enzyme combinations. Since the embeddings are $L_2$-normalized, the dot product $\vec{\mathbf{f}}_R(r;\vec{\boldsymbol{\theta}}) \cdot \vec{\mathbf{f}}_P(p;\vec{\boldsymbol{\theta}})$ is equivalent to the cosine similarity.

Assuming independent and identically distributed samples, the likelihood of observing the annotated set $\mathcal{A}$ is

$$P(\mathcal{A}|\vec{\boldsymbol{\theta}}) = \prod_{(r,p)\in\mathcal{A}} P(r, p|\vec{\boldsymbol{\theta}}). \tag{2}$$

*

We train the model by minimizing the negative log-likelihood, defined as the MLNCE loss:

$$\begin{aligned}\mathcal{L}_{\text{MLNCE}}(\vec{\boldsymbol{\theta}};\mathcal{A}) &= -\frac{1}{|\mathcal{A}|}\sum_{(r,p)\in\mathcal{A}}\log P(r,p|\vec{\boldsymbol{\theta}}) \\ &= -\frac{1}{|\mathcal{A}|}\sum_{(r,p)\in\mathcal{A}}\Big[\beta\vec{\mathbf{f}}_R(r;\vec{\boldsymbol{\theta}})\cdot\vec{\mathbf{f}}_P(p;\vec{\boldsymbol{\theta}}) - \log Z(\vec{\boldsymbol{\theta}})\Big] \\ &= \log Z(\vec{\boldsymbol{\theta}}) - \frac{1}{|\mathcal{A}|}\sum_{(r,p)\in\mathcal{A}}\beta\vec{\mathbf{f}}_R(r;\vec{\boldsymbol{\theta}})\cdot\vec{\mathbf{f}}_P(p;\vec{\boldsymbol{\theta}}).\end{aligned} \tag{3}$$

Calculating $Z(\vec{\boldsymbol{\theta}})$ over all possible pairs is computationally intractable. Therefore, we approximate the partition function using only the reactions and enzymes present within a single training batch. We partition $\mathcal{A}$ into $N_B$ batches such that $\bigcup_{b=1}^{N_B}\mathcal{A}_b = \mathcal{A}$, with corresponding entity subsets $\mathcal{R}_b$ and $\mathcal{P}_b$. The approximated batch partition function is

$$Z_b(\vec{\boldsymbol{\theta}}) = \sum_{(r',p')\in\mathcal{R}_b\times\mathcal{P}_b} e^{\beta\vec{\mathbf{f}}_R(r';\vec{\boldsymbol{\theta}})\cdot\vec{\mathbf{f}}_P(p';\vec{\boldsymbol{\theta}})}. \tag{4}$$

The final batch-approximated loss becomes

$$\mathcal{L}(\vec{\boldsymbol{\theta}};\mathcal{A}) \approx \sum_{b=1}^{N_B}\left[\log Z_b(\vec{\boldsymbol{\theta}}) - \frac{1}{|\mathcal{A}_b|}\sum_{(r,p)\in\mathcal{A}_b}\beta\vec{\mathbf{f}}_R(r;\vec{\boldsymbol{\theta}})\cdot\vec{\mathbf{f}}_P(p;\vec{\boldsymbol{\theta}})\right]. \tag{5}$$

This approach differs from the standard InfoNCE loss used in contrastive learning, which maximizes the conditional probability of one modality given the other — identifying the correct enzyme for a specific reaction (or vice-versa)–—rather than the joint probability of the pair [18]. The two possible conditional probabilities are defined as

$$P(p|r,\vec{\boldsymbol{\theta}}) = \frac{e^{\beta\vec{\mathbf{f}}_R(r;\vec{\boldsymbol{\theta}})\cdot\vec{\mathbf{f}}_P(p;\vec{\boldsymbol{\theta}})}}{\sum_{p'\in\mathcal{P}} e^{\beta\vec{\mathbf{f}}_R(r;\vec{\boldsymbol{\theta}})\cdot\vec{\mathbf{f}}_P(p';\vec{\boldsymbol{\theta}})}} = \frac{e^{\beta\vec{\mathbf{f}}_R(r;\vec{\boldsymbol{\theta}})\cdot\vec{\mathbf{f}}_P(p;\vec{\boldsymbol{\theta}})}}{Z_r(\vec{\boldsymbol{\theta}})}, \tag{6}$$

$$P(r|p,\vec{\boldsymbol{\theta}}) = \frac{e^{\beta\vec{\mathbf{f}}_R(r;\vec{\boldsymbol{\theta}})\cdot\vec{\mathbf{f}}_P(p;\vec{\boldsymbol{\theta}})}}{\sum_{r'\in\mathcal{R}} e^{\beta\vec{\mathbf{f}}_R(r';\vec{\boldsymbol{\theta}})\cdot\vec{\mathbf{f}}_P(p;\vec{\boldsymbol{\theta}})}} = \frac{e^{\beta\vec{\mathbf{f}}_R(r;\vec{\boldsymbol{\theta}})\cdot\vec{\mathbf{f}}_P(p;\vec{\boldsymbol{\theta}})}}{Z_p(\vec{\boldsymbol{\theta}})}, \tag{7}$$

with corresponding loss function

$$\begin{aligned}\mathcal{L}_{\text{InfoNCE}}(\vec{\boldsymbol{\theta}};\mathcal{A}) &= -\frac{1}{|\mathcal{A}|}\sum_{(r,p)\in\mathcal{A}}\log P(p|r,\vec{\boldsymbol{\theta}}), \\ \mathcal{L}_{\text{InfoNCE}}(\vec{\boldsymbol{\theta}};\mathcal{A}) &= -\frac{1}{|\mathcal{A}|}\sum_{(r,p)\in\mathcal{A}}\log P(r|p,\vec{\boldsymbol{\theta}}).\end{aligned} \tag{8}$$

CLIP utilizes a symmetrized loss that averages these two conditional views,

$$\mathcal{L}_{\text{CLIP}}(\vec{\boldsymbol{\theta}};\mathcal{A}) = -\frac{1}{|\mathcal{A}|}\sum_{(r,p)\in\mathcal{A}}\frac{1}{2}\Big[\log P(r|p,\vec{\boldsymbol{\theta}}) + \log P(p|r,\vec{\boldsymbol{\theta}})\Big]. \tag{9}$$

In practice, we find that the MLNCE loss is less susceptible to overfitting. We hypothesize that MLNCE improves performance over InfoNCE/CLIP by contrasting more negative examples against each positive pair. Consider a batch containing $N$ unique reaction-enzyme pairs.

- **InfoNCE/CLIP (Linear Scaling):** These losses normalize over a single row or column of the similarity matrix. For a given reaction $r$, the model contrasts the true enzyme $p$ against $N-1$ other enzymes in the batch. Thus, the number of negative samples scales linearly with batch size ($\propto N$).
- **MLNCE (Quadratic Scaling):** The MLNCE partition function $Z_b$ sums over the entire $N \times N$ matrix of all possible cross-pairs in the batch. Every unannotated combination in the batch contributes to the denominator, meaning the model implicitly contrasts the positive pair against $N^2 - 1$ negative pairs.

Crucially, this quadratic increase in negative samples does not significantly increase computational cost. Since both methods require computing the full $N \times N$ pairwise similarity matrix, MLNCE simply utilizes the off-diagonal terms that InfoNCE ignores. However, in-batch negative sampling in all cases comes with a trade-off: it relies on the single batch partition function effectively approximating the global partition function. Small batches may yield high variance in this approximation, explaining our empirical observation that MLNCE benefits significantly from larger batch sizes.

#### PROTEIN LANGUAGE MODEL COMPUTE COST AND FINETUNING ANALYSIS

A key consideration in model development is the trade-off between performance and computational efficiency. Large transformer-based models, like protein language models (pLMs), can demand orders of magnitude more compute resources for training or fine-tuning compared to smaller architectures such as multilayer perceptrons (MLPs). We therefore analyzed the relative performance of our different protein embedding strategies against their estimated computational cost, measured in floating point operations (FLOPs).

##### Dataset Preparation

To compare different enzyme embedding schemes, we used a subset of our development dataset. Sequences were clustered at 80% similarity using MMseqs2 [21] to reduce redundancy. This dataset was subdivided into a fixed 80%/20% training/validation split based directly on reactions, rather than reaction fingerprint clusters. For models utilizing structural information, we downloaded 3D coordinates from the AlphaFold Database [9, 23].

##### Model Training Protocol

All 650M parameter model training utilized 4 NVIDIA A100 GPUs with DeepSpeed ZeRO-2 for distributed training and memory management. This approach balances memory savings with communication volume, a critical factor during full fine-tuning. Finally, models were evaluated on the validation set using all proteins from the development dataset (clustered at 80% sequence similarity) as the screening set.

##### Protein Embedding Strategies

We compared two main paradigms for using pLMs to generate protein embeddings:

1. **Pre-computed Embeddings:** This approach involves a one-time process where a pre-trained pLM generates fixed embeddings for the entire protein dataset. These static embeddings are then reused across all training epochs. While the initial generation incurs a significant one-time cost, this is amortized over the training duration. We tested ESM-2 (650M), ProtT5-XL-U50 (ProtT5 3B), and ESM-2 (3B). These models were trained for 100 epochs using batch sizes of 32 (matching the live models) and 16,384 (reference).

2. **Live pLM Integration:** This method keeps the pLM weights loaded in memory during training, allowing gradients to propagate through the model. This enables fine-tuning but substantially increases the computational cost per step. We investigated two architectures within this paradigm, using ESM-2 (650M):

   - **Mean-Pooling:** Sequence embeddings were generated by mean-pooling the final hidden states of all tokens, which were then passed to a final MLP layer.
   - **Structure-Biased Attention Pooling:** To incorporate 3D structural information, we augmented the model with a structure-aware attention layer inserted before the final pooling step. This layer uses a hidden dimension of 1280, a single attention head, and a 4× expanded feed-forward dimension of 5120. The self-attention mechanism is biased by the inverse pairwise Euclidean distances between residues (calculated from AlphaFold structures), encouraging attention between spatially proximal residues while maintaining SE(3) invariance.

   Training durations were tailored to each live model configuration to balance convergence and computational cost:

   - **Mean-Pooling (Frozen Base):** Trained for 45 epochs.
   - **Mean-Pooling (Fine-Tuned):** Trained for 10 epochs.
   - **Structure-Biased Attention (Frozen Base):** Trained for 60 epochs.
   - **Structure-Biased Attention (Fine-Tuned):** Trained for 60 epochs.

##### Computational Cost Analysis

Larger models inherently require more FLOPs per training step and often necessitate longer training durations and more powerful hardware, impacting both financial costs and the speed of experimental iteration. Furthermore, the considerable memory footprint of models like ESM-2 (650M) often compels the use of smaller batch sizes compared to models trained on pre-computed embeddings. For instance, our ESM-based models were limited to a batch size of 32. In contrast, models using pre-computed embeddings could potentially leverage much larger batch sizes (e.g., 16,384, as used for a reference model comparison), which can also influence training dynamics and final performance.

Furthermore, the contrastive loss function has a unique complication with respect to batch size. Typically, when data points are independent and identically distributed (i.i.d.), a desired global batch size can easily be achieved through combining batches via gradient accumulation and data parallel training despite small local mini-batch sizes. However, this averaging cannot be employed so trivially with standard contrastive losses that employ random negative sampling, as all pairs in a batch contribute to the denominator of each loss term. An online-MLNCE loss function, drawing from the online-softmax implementations, is an area for further work.

Given these trade-offs, a central question emerges: Does the potential performance gain from unfreezing the pLM, or from adding architectural modifications like a structure-aware layer, justify the dramatically increased computational expense? To address whether the computational expense of unfreezing the pLM or adding structural layers is justified, we compared the performance-per-FLOP across configurations. By translating the training epochs for each model into an estimated total FLOP count, we establish a fairer comparison based on a fixed computational budget. The results, presented in Fig. S2, illustrate the top-1, top-10, and top-100 hit rates on the validation set plotted against cumulative FLOPs, enabling a direct comparison of efficiency across strategies.

Unsurprisingly, models using pre-computed embeddings achieve the best final performance for the lowest computational cost. When comparing the live ESM-2 models, the frozen configurations generally yield better performance per FLOP than their unfrozen counterparts for the more challenging top-1 and top-10 hit rate tasks, although both significantly lag behind the pre-computed embedding approach in overall efficiency. However, the unfrozen models demonstrate competitive performance for the top-100 hit rate metric relative to their FLOP cost. Ultimately, the performance of all live ESM-2 models (frozen or unfrozen) appears constrained by the small batch size (32) necessitated by memory limitations, compared to the much larger batch size (16k) feasible with pre-computed embeddings.

#### CLIPZYME BENCHMARKING AND INFERENCE COMPARISON

##### Computational Benchmark

To compare our model against CLIPZyme [14], we utilized the specific version of the EnzymeMap dataset provided by the authors [15]. This dataset contains reaction-enzyme pairs and a separate screening set of unpaired enzymes. As the original data splits were not explicitly included, we used the authors' provided code (`https://github.com/pgmikhael/clipzyme`) to replicate their training, development (dev), and test splits as closely as possible.

During this replication, we encountered several discrepancies regarding the statistics reported in the original publication. Despite these differences, we determined that the comparison remains valid. The discrepancies are detailed below:

- **Reaction Splits:** The reported train/dev/test reaction counts (12,629 / 2,669 / 1,554) differed slightly from our observed counts (12,603 / 2,661 / 1,551).
- **Screening Set Size:** The reported size (260,197) differed from the observed size (261,907).
- **Empty Sequences:** We identified 72 empty sequences within the screening set, likely due to deprecated UniProt entries. Removing these reduced the effective screening set size to 261,835 unique sequences. Some of these missing/empty sequences also affected the paired reaction data, resulting in reduced train/dev/test pair counts (from 34,427 / 7,287 / 4,642 to 34,265 / 7,285 / 4,638) and reaction counts (12,569 / 2,661 / 1,551).
- **Data Contamination:** Crucially, we note that the dataset was split based on reaction-enzyme pairs, not solely on reactions. This resulted in data leakage, with 9 training reactions appearing in the dev set and 30 training reactions appearing in the test set.

For the benchmarking protocol (Table S5), we utilized the dataset described above with the standard 80%/10%/10% train/dev/test split. Models were trained for 30 epochs and evaluated on the test split using the screening set provided within the CLIPZyme distribution.

##### Performance on Experimental Case Studies

We further evaluated CLIPZyme's ability to recover the experimentally verified predictions generated by *Horizyn-1*. This analysis focused on two distinct tasks:

First, for **enzyme discovery tasks** (the orphan reactions from Fig. 2 and transaminase reactions for non-canonical amino acid synthesis from Fig. 3), we queried each reaction against the CLIPZyme enzyme screening set. This analysis was constrained by the coverage of the CLIPZyme screening set:

- For the orphan reactions, the native enzymes were absent from the screening set; therefore, *E. coli* GabT was used as the closest available homolog for evaluation.
- For the transaminase reactions, only three of the relevant enzymes were present in the screening set.

Second, for the **functional annotation** of uncharacterized *Thermus aquaticus* enzymes, we queried the sequences against the full CLIPZyme reaction dataset. In this instance, both ground-truth target reactions were confirmed to be present within the dataset.

In all evaluated cases, CLIPZyme failed to rank the targets sufficiently high for successful retrieval. A summary of this analysis is provided in Table S6, Table S7, and Table S8.

#### PROTEIN FUNCTION ANNOTATION COMPARISON

To provide a comparative baseline for the *Horizyn-1* annotations of uncharacterized *Thermus aquaticus* enzymes (see Fig. 2, we evaluated several popular alternative protein function prediction methods.

First, we employed alignment and structure-based search tools—specifically MMseqs2 [21], BLASTp [3], and Foldseek [22]—using default parameters. For each tool, we performed a search against our inference screening set of ∼7 million enzymes. The retrieved enzymes were mapped to their corresponding reactions in the inference dataset. To score a specific reaction, we assigned it the maximum bitscore observed among all enzymes mapped to that reaction. Reactions were then ranked in descending order based on these scores.

Additionally, we utilized the CLEAN webserver (`https://clean.platform.ibiofoundry.illinois.edu/configuration`) [25] to retrieve predicted EC numbers directly from the input sequences. For details regarding the CLIPZyme baseline, see CLIPZyme Benchmarking and Inference Comparison. The results of this comparative analysis are summarized in Table S7.

#### HARD NEGATIVE SAMPLING ANALYSIS

To evaluate the efficacy of our random in-batch sampling strategy (see Contrastive Learning and Negative Sampling), we compared it against a baseline model trained using hard negative mining. Hard negatives are typically defined as unannotated pairs that are difficult to distinguish from positive pairs due to high similarity. We constructed these hard negatives based on EC numbers, similar to the protocol in CLEAN [25].

Prior to training, for every positive reaction-enzyme pair in the training split of the development dataset, we sampled specific unannotated hard negative pairs where the reaction and enzyme shared the same EC number up to the third (EC3) or fourth (EC4) level digits. This ensured the negative examples were functionally similar to true reaction-enzyme pairs but not explicitly annotated in the dataset. Unlike our random in-batch sampling approach, which generates negatives dynamically, these hard negatives were constructed offline. The model was then trained to distinguish positive pairs strictly from these pre-selected hard negatives, without the inclusion of random background noise.

This setup requires a modification to the probability definition in (1) used for the standard MLNCE loss. Let $\mathcal{A}$ be the set of annotated positive pairs and $\mathcal{N}$ be the set of precomputed hard negative pairs. We define the Hard Negative loss as

$$\mathcal{L}_{\text{Hard}} = -\frac{1}{|\mathcal{A}|} \sum_{(r,p)\in\mathcal{A}} \log P(r,p|\vec{\boldsymbol{\theta}}) \tag{10}$$

using the modified probability

$$P(r,p|\vec{\boldsymbol{\theta}}) = \frac{e^{\beta \vec{\mathbf{f}}_R(r;\vec{\boldsymbol{\theta}})\cdot\vec{\mathbf{f}}_P(p;\vec{\boldsymbol{\theta}})}}{\sum_{(r',p')\in\mathcal{A}\cup\mathcal{N}} e^{\beta \vec{\mathbf{f}}_R(r';\vec{\boldsymbol{\theta}})\cdot\vec{\mathbf{f}}_P(p';\vec{\boldsymbol{\theta}})}}. \tag{11}$$

Note that the denominator no longer sums over all possible pairs in the dataset, but instead restricted to the annotated positive and precomputed hard negative pairs.

In practice, we pool the annotated and hard negative pairs into a unified dataset $\mathcal{D} = \mathcal{A} \cup \mathcal{N}$. We then randomly partition $\mathcal{D}$ into a set of batches $\bigcup_{b=1}^{N_B} \mathcal{D}_b = \mathcal{D}$, where each batch $\mathcal{D}_b$ contains a random mixture of positive and negative pairs. The probability for a pair within a given batch is approximated by normalizing over all samples present in that batch, with partition function

$$Z_b(\vec{\boldsymbol{\theta}}) = \sum_{(r',p')\in\mathcal{D}_b} e^{\beta\vec{\mathbf{f}}_R(r';\vec{\boldsymbol{\theta}})\cdot\vec{\mathbf{f}}_P(p';\vec{\boldsymbol{\theta}})}. \tag{12}$$

and batch-approximated loss

$$\mathcal{L}(\vec{\boldsymbol{\theta}};\mathcal{A}) \approx \sum_{b=1}^{N_B}\left[\log Z_b(\vec{\boldsymbol{\theta}}) - \frac{1}{|\mathcal{D}_b|}\sum_{(r,p)\in\mathcal{D}_b}\beta\vec{\mathbf{f}}_R(r;\vec{\boldsymbol{\theta}})\cdot\vec{\mathbf{f}}_P(p;\vec{\boldsymbol{\theta}})\right]. \tag{13}$$

The results of this analysis are demonstrated in Table S4. For hard EC4 sampling, we utilized all possible negatives ( 300k), while for EC3, we randomly subsampled varying numbers of hard negatives.

#### MISSING PAIR ANALYSIS

Due to enzyme promiscuity, curated datasets rarely include all possible reaction-enzyme pairs. Consequently, the model risks learning the probability of a pair being annotated rather than the probability of an enzyme catalyzing a particular reaction, potentially treating unannotated true positives as negatives. To evaluate the model's robustness to this missing information, we simulated an incomplete data scenario by randomly subsetting the development dataset to 80% of the original pairs. Because reactions and proteins often participate in multiple independent pairs, this generated a significant number of cases where both the reaction and enzyme were included in the training set (via other associations), but the specific pair was excluded and therefore treated as a negative. We trained a model on this subset using the standard development protocol. We then classified all possible reaction-enzyme combinations into three categories for evaluation: (i) included positive pairs, which were explicitly present in the 80% training subset; (ii) missing positive pairs, which were valid pairs in the development dataset but excluded from the training subset; and (iii) negative pairs, where the protein and reaction are not associated. Results are depicted in Fig. S3.

#### SEQUENCE CLUSTERING ANALYSIS

To investigate the effects of sequence similarity clustering on model performance, we clustered the training split of the development dataset and evaluated the resulting models on the test split. Clustering was performed using MMseqs2 with varying sequence identity thresholds (refer to the Inference Dataset section for specific command arguments). For the 100% sequence identity threshold, we employed exact string matching to identify duplicates. The results of this analysis are summarized in Fig. S4.

#### REACTION SIMILARITY CLUSTERING ANALYSIS

The strategy for splitting data into training, validation, and test sets is crucial for obtaining realistic performance estimates. We advocate for splitting datasets at the reaction level, rather than, for example, splitting on reaction-enzyme pairs or enzymes. This approach directly reflects the real-world use case where models must generalize to entirely new, unseen classes of reactions.

A key challenge in reaction splitting is avoiding data contamination, where reactions highly similar to those in the training set appear in the validation or test sets. Such leakage can lead to inflated performance metrics that do not translate to real-world utility. To mitigate this, we cluster reactions based on their similarity and ensure that all reactions within a cluster are assigned to the same data split.

Several methods have been employed historically for assessing reaction similarity for clustering. Template-based approaches, such as those used in methods like CLIPZyme [14], attempt to identify conserved reaction patterns. However, templates may not be available for all reaction types, and a single reaction can sometimes be assigned multiple templates, leading to ambiguity. Another approach, employed by methods like Reactzyme [7], uses fuzzy string matching on reaction SMILES strings. While more general, this method can be susceptible to inconsistencies arising from the canonicalization process of SMILES representations.

Given these limitations, we used a historically standard and robust method: representing reactions with structural reaction fingerprints that concatenate extended-connectivity fingerprints (ECFP6, also known as Morgan fingerprints) [17, 19] of the reactants and products and quantifying their similarity using the Tanimoto (Jaccard) coefficient (see Structural Fingerprint Clustering Protocol). This approach is broadly applicable to diverse reaction types and exhibits stable behavior.

##### Template Clustering Protocol

For comparison purposes, we also constructed splits using reaction templates. We generated atom mappings between reactants and products for all reactions using the RXNMapper transformer model [20]. Subsequently, we constructed the condensed graph of reaction (also referred to as "imaginary transition states") [5, 24], retaining only the atoms and bonds involved in bonding changes. The retained substructures of the reactants and products were saved as reaction templates in SMARTS notation. Due to limitations in atom mapping, valid reaction templates could only be constructed for 10,473 of the 11,932 Rhea reactions.

##### Split Difficulty Analysis

To understand the model's generalization capabilities (Fig. S12), we systematically analyzed the validation error as a function of the similarity threshold used during data splitting. We used the training portion of the development dataset (see Development Model Training and Evaluation), subdividing it into approximate 80%/20% training/validation splits using different random seeds for each replicate.

These splits were generated based on either:

- **Fingerprint Clusters:** Using the Tanimoto similarity threshold defined in the main text (see *Structural Fingerprint Clustering Protocol*).
- **Template Clusters:** Using the templates described above. Reactions that could not be mapped to a valid template were assigned to the training set; the remaining reactions were then randomly divided to meet the 80%/20% ratio.

All models for this analysis were trained for 100 epochs and evaluated using the corresponding validation set. The entire protein set from the full development dataset was used as the screening set for all reaction queries.

#### ENZYME PROMISCUITY BENCHMARK

Enzyme promiscuity with respect to reaction substrates was evaluated on a collection of 6 published enzyme-substrate reactivity datasets by S. Goldman (`https://github.com/samgoldman97/enzyme-datasets`) (Fig. S15). Each dataset contained the full tableau of enzyme-substrate pairs with positive (reactive) and negative (non-reactive) annotations. The following datasets were included in this study;

1. Aminotransferases (25 enzymes, 18 substrates) [12]
2. DUF849 (beta-keto acid cleavage enzymes, BKACE, 161 enzymes, 17 substrates) [1]
3. Esterases (146 enzymes, 96 substrates) [13]
4. Nitrilases (18 enzymes, 38 substrates) [2]
5. OleA thiolases (73 enzymes, 15 substrates) [16]
6. Phosphatases (218 enzymes, 165 substrates) [8]

Quantitative activity measurements were available for three of these datasets, defined as follows:

1. Aminotransferases: specific activity (U/mg) (U = enzyme activity unit, $\mu$mol/min)
2. OleA thiolases: maximum slope of the fluorescence signal at $\lambda = 410$ nm ($p$-nitrophenolate)

3. Phosphatases: absorbance at $\lambda = 650$ nm

Duplicate substrate records were removed. In the phosphatase dataset, incorrect substrate SMILES strings were replaced by SMILES retrieved from PubChem [11] by substrate name. Reaction SMILES strings were generated from the substrate SMILES by applying reaction-specific SMARTS templates, shown in Table S9. When multiple alternative products were possible, all possibilities were generated, and the highest Horizyn-1 pair score was used in downstream tasks. The final dataset sizes are given in Table S10.

---

[11] S. Kim, J. Chen, T. Cheng, A. Gindulyte, J. He, S. He, Q. Li, B. A. Shoemaker, P. A. Thiessen, B. Yu, L. Zaslavsky, J. Zhang, and E. E. Bolton. PubChem 2025 update. *Nucleic Acids Research*, 53(D1):D1516–D1525, 2025.

[12] T. Li, X. Cui, Y. Cui, J. Sun, Y. Chen, T. Zhu, C. Li, R. Li, and B. Wu. Exploration of transaminase diversity for the oxidative conversion of natural amino acids into 2-ketoacids and high-value chemicals. *ACS Catalysis*, 10(14):7950–7957, July 2020.

[13] M. Martínez-Martínez, C. Coscolín, G. Santiago, J. Chow, P. J. Stogios, R. Bargiela, C. Gertler, J. Navarro-Fernández, A. Bollinger, S. Thies, C. Méndez-García, A. Popovic, G. Brown, T. N. Chernikova, A. García-Moyano, G. E. K. Bjerga, P. Pérez-García, T. Hai, M. V. Del Pozo, R. Stokke, I. H. Steen, H. Cui, X. Xu, B. P. Nocek, M. Alcaide, M. Distaso, V. Mesa, A. I. Peláez, J. Sánchez, P. C. F. Buchholz, J. Pleiss, A. Fernández-Guerra, F. O. Glöckner, O. V. Golyshina, M. M. Yakimov, A. Savchenko, K.-E. Jaeger, A. F. Yakunin, W. R. Streit, P. N. Golyshin, V. Guallar, M. Ferrer, and The INMARE Consortium. Determinants and prediction of esterase substrate promiscuity patterns. *ACS Chemical Biology*, 13(1):225–234, Jan. 2018.

[14] P. Mikhael, I. Chinn, and R. Barzilay. CLIPZyme: Reaction-conditioned virtual screening of enzymes. In R. Salakhutdinov, Z. Kolter, K. Heller, A. Weller, N. Oliver, J. Scarlett, and F. Berkenkamp, editors, *Proceedings of the 41st International Conference on Machine Learning (ICML)*, pages 35647–35663. PMLR, 2024.

[15] P. Mikhael, I. Chinn, and R. Barzilay. Data from "CLIPZyme: Reaction-conditioned virtual screening of enzymes", 2024. Zenodo. Available at `https://doi.org/10.5281/zenodo.15161343`. Deposited 5 April 2025.

[16] S. L. Robinson, M. D. Smith, J. E. Richman, K. G. Aukema, and L. P. Wackett. Machine learning-based prediction of activity and substrate specificity for OleA enzymes in the thiolase superfamily. *Synthetic Biology*, 5(1), Jan. 2020.

[17] D. Rogers and M. Hahn. Extended-connectivity fingerprints. *Journal of Chemical Information and Modeling*, 50(5):742–754, May 2010.

[18] E. Rusak, P. Reizinger, A. Juhos, O. Bringmann, R. S. Zimmermann, and W. Brendel. InfoNCE: Identifying the gap between theory and practice. In Y. Li, S. Mandt, S. Agrawal, and E. Khan, editors, *Proceedings of The 28th International*

*Conference on Artificial Intelligence and Statistics (AISTATS)*, pages 4159–4167. PMLR, 2025.

[19] N. Schneider, D. M. Lowe, R. A. Sayle, and G. A. Landrum. Development of a novel fingerprint for chemical reactions and its application to large-scale reaction classification and similarity. *Journal of Chemical Information and Modeling*, 55(1):39–53, Jan. 2015.

[20] P. Schwaller, B. Hoover, J.-L. Reymond, H. Strobelt, and T. Laino. Extraction of organic chemistry grammar from unsupervised learning of chemical reactions. *Science Advances*, 7(15):eabe4166, Apr. 2021.

[21] M. Steinegger and J. Söding. MMseqs2 enables sensitive protein sequence searching for the analysis of massive data sets. *Nature Biotechnology*, 35(11):1026–1028, Oct. 2017.

[22] M. van Kempen, S. S. Kim, C. Tumescheit, M. Mirdita, J. Lee, C. L. M. Gilchrist, J. Söding, and M. Steinegger. Fast and accurate protein structure search with Foldseek. *Nature Biotechnololgy*, 42(2):243–246, Feb. 2024.

[23] M. Varadi, D. Bertoni, P. Magana, U. Paramval, I. Pidruchna, M. Radhakrishnan, M. Tsenkov, S. Nair, M. Mirdita, J. Yeo, O. Kovalevskiy, K. Tunyasuvunakool, A. Laydon, A. Žídek, H. Tomlinson, D. Hariharan, J. Abrahamson, T. Green, J. Jumper, E. Birney, M. Steinegger, D. Hassabis, and S. Velankar. AlphaFold protein structure database in 2024: providing structure coverage for over 214 million protein sequences. *Nucleic Acids Research*, 52(D1):D368–D375, Jan. 2024.

[24] A. Varnek, D. Fourches, F. Hoonakker, and V. P. Solov'ev. Substructural fragments: an universal language to encode reactions, molecular and supramolecular structures. *Journal of Computer-Aided Molecular Design*, 19(9-10):693–703, Sept. 2005.

[25] T. Yu, H. Cui, J. C. Li, Y. Luo, G. Jiang, and H. Zhao. Enzyme function prediction using contrastive learning. *Science*, 379(6639):1358–1363, Mar. 2023.

| $k$ | **Top-$k$ Hit Rate** | **Prec@$k$** | **Recall@$k$** | **F1@$k$** |
|---|---|---|---|---|
| 1 | 36.2 | 36.2 | 14.5 | 17.4 |
| 10 | 57.1 | 20.9 | 43.3 | 21.1 |
| 100 | 76.7 | 7.1 | 68.5 | 9.3 |
| 1000 | 92.1 | 1.9 | 86.9 | 3.1 |
| $R$ | 49.3 | 36.9 | 36.9 | 36.9 |

TABLE S1. **Precision, recall, and F1 score at different top-$k$ thresholds.** Detailed retrieval metrics at varying top-$k$ thresholds for the development model (Fig. 1D) on the development test set. In the final row, $k$ is set dynamically to the total number of true positives ($R$) for each specific query (i.e., Prec@$R$ = $R$-precision).

| **EC Class** | **Training Reactions** | **Test Reactions** | **Top-1 Hit Rate** | **Top-10 Hit Rate** | **Top-100 Hit Rate** | **Top-1000 Hit Rate** | **$R$-Precision** | **Average Precision** |
|---|---|---|---|---|---|---|---|---|
| 1.-.-.- | 3508 (33%) | 321 (32%) | 41.4 | 62.6 | 83.8 | 94.8 | 40.3 | 46.4 |
| 2.-.-.- | 3507 (33%) | 314 (31%) | 35.5 | 58.6 | 79.3 | 95.0 | 37.9 | 42.7 |
| 3.-.-.- | 1838 (17%) | 188 (19%) | 39.9 | 63.6 | 74.6 | 91.4 | 42.3 | 45.6 |
| 4.-.-.- | 1096 (10%) | 151 (15%) | 21.6 | 45.1 | 72.4 | 90.7 | 26.1 | 29.2 |
| 5.-.-.- | 446 (4%) | 71 (7%) | 18.7 | 44.6 | 69.9 | 86.7 | 20.6 | 25.6 |
| 6.-.-.- | 413 (4%) | 37 (4%) | 25.6 | 41.1 | 55.6 | 78.9 | 23.1 | 28.1 |
| 7.-.-.- | 194 (2%) | 5 (<1%) | 40.0 | 60.0 | 80.0 | 100.0 | 43.6 | 47.9 |
| N/A | 932 (9%) | 59 (6%) | 55.1 | 67.8 | 78.0 | 93.2 | 50.4 | 55.9 |

TABLE S2. **Performance of Horizyn-1 by EC number.** Distribution of top-level EC classes in the train and test splits of the development dataset (Rhea/Swiss-Prot) and corresponding performance of the development model (Fig. 1D) for each class. No correlation is observed between model performance within an EC class and the number of training reactions within the class. Parentheses in the reaction columns indicate the percentage of the total dataset represented by that class.

| **Model / EC Level** | **Top-1 Hit Rate** | **Top-10 Hit Rate** | **Top-100 Hit Rate** | **Top-1000 Hit Rate** | **$R$-Precision** | **Average Precision** |
|---|---|---|---|---|---|---|
| EC 1 (a.-.-.-) | 0.1 | 0.7 | 5.2 | 20.6 | 0.1 | 0.1 |
| EC 2 (a.b.-.-) | 7.1 | 4.5 | 17.7 | 51.3 | 0.7 | 0.9 |
| EC 3 (a.b.c.-) | 2.2 | 10.3 | 32.4 | 69.9 | 2.1 | 2.9 |
| EC 4 (a.b.c.d) | 55.5 | 74.7 | 85.4 | 90.3 | 55.5 | 58.3 |
| *Horizyn-1* | 36.2 | 57.1 | 76.7 | 92.1 | 36.9 | 42.1 |
| *Horizyn-1* + EC 1 (a.-.-.-) | 37.3 | 59.6 | 80.7 | 95.0 | 38.7 | 44.0 |
| *Horizyn-1* + EC 2 (a.b.-.-) | 38.8 | 62.6 | 84.6 | 97.0 | 40.8 | 46.3 |
| *Horizyn-1* + EC 3 (a.b.c.-) | 41.1 | 65.3 | 86.7 | 97.9 | 43.3 | 48.9 |
| *Horizyn-1* + EC 4 (a.b.c.d) | 77.1 | 92.3 | 97.2 | 99.3 | 77.5 | 80.4 |

TABLE S3. **Performance conditioned on EC numbers.** Development model performance (Fig. 1D) compared to rankings based on partial EC numbers and a composite score combining both methods. *Horizyn-1* scores are computed using the development model (Fig. 1D). For the composite method, enzymes are prioritized by EC number matches, with *Horizyn-1* scores used to rank enzymes within those groups. *Horizyn-1* outperforms a ranking based solely on EC numbers up to the third digit, while combing *Horizyn-1* with fourth digit EC numbers dramatically boosts performance. Full analysis details can be found in the EC Conditioning Analysis section of the Methods.

| Negative Sampling Strategy | Number of Negatives | Training Time | Top-1 Hit Rate | Top-10 Hit Rate | Top-100 Hit Rate | Top-1000 Hit Rate | $R$-Precision | Average Precision |
|---|---|---|---|---|---|---|---|---|
| Hard EC4 | ∼300K | 2h 16m | 9.5 | 14.3 | 21.4 | 38.3 | 8.0 | 8.8 |
| Hard EC3 (1M) | 1M | 5h 1m | 27.6 | 48.0 | 64.2 | 80.9 | 28.2 | 32.2 |
| Hard EC3 (2M) | 2M | 8h 38m | 30.7 | 49.4 | 62.8 | 77.6 | 30.4 | 34.4 |
| Hard EC3 (4M) | 4M | 15h 20m | 29.6 | 47.8 | 61.7 | 76.1 | 29.6 | 33.4 |
| Random | ∼2B | 1h 24m | 36.2 | 57.1 | 76.7 | 92.1 | 36.9 | 42.1 |

TABLE S4. **Performance comparison of hard negative vs. random sampling.** In-batch random negative sampling is evaluated against baselines using pre-computed hard negatives based on EC number similarity (EC3/EC4). The random sampling approach yielded significantly higher retrieval metrics. This improvement is attributed to the computational efficiency of in-batch sampling, which allows for a three orders of magnitude increase in the number of negative samples per training step compared to the hard mining baseline despite a significantly reduced training time (see Hard Negative Sampling Analysis for details).

| Model / Source | Reaction Embeddings | Enzyme Embeddings | BEDROC ($\alpha = 20$) | BEDROC ($\alpha = 85$) | EF (5%) | EF (10%) |
|---|---|---|---|---|---|---|
| *Horizyn-1* | ECFP6/DRFP | ProtT5 | 66.3 | 48.5 | 14.7 | 8.3 |
| | ECFP6/DRFP | ESM-2 (3B) | 59.8 | 41.6 | 13.5 | 7.6 |
| | ECFP6/DRFP | ESM-2 (650M) | 57.8 | 42.4 | 12.7 | 7.3 |
| Ref. [14] | PTS-GNN | ESM-2 (650M) | 29.4 | 17.8 | 6.6 | 4.2 |
| CLIPZyme [14] | PTS-GNN | ESM-2 (650M) + EGNN | 63.0 | 44.7 | 14.1 | 8.1 |

TABLE S5. **Comparison of *Horizyn-1* and *CLIPZyme* on the EnzymeMap benchmark.** Performance of *Horizyn-1* (trained on the EnzymeMap training split) and CLIPZyme variants was assessed using early recognition metrics. Specifically, we report the Boltzmann-Enhanced Discrimination of ROC (BEDROC) with $\alpha = 20$ and 85, and the Enrichment Factor (EF) at the top 5% and 10% of the ranked list. Full definitions for these metrics are located in the Model Evaluation section. *Horizyn-1* variants combine ECFP6/DRFP reaction fingerprints with different protein language models [ProtT5, ESM-2 (3B), and ESM-2 (650M)]. Data for the CLIPZyme variants are reproduced from [14]; these models use a pseudo-transition state graph neural network (PTS-GNN) for reaction embeddings and ESM-2 (650M) for protein embeddings, with and without a structure-aware EGNN module.

| Reaction | *Horizyn-1* | CLIPZyme |
|---|---|---|
| 6-Aminohexanoate (RHEA:58200) | 1 | 3194 |
| $\beta$-Alanine (RHEA:30699) | 379 | 1576 |
| Putrescine (RHEA:23816) | 30 | 80 |

TABLE S6. **Comparison of *Horizyn-1* and CLIPZyme on enzyme discovery for orphan reactions.** The predicted rank of *E. coli* GabT (P22256) (see Fig. 2) is listed within each model's respective screening set. *Horizyn-1*'s screening set consists of ∼7 million enzymes, while CLIPZyme considers ∼260,000. *Horizyn-1* Despite a substantially larger screening set, *Horizyn-1* consistently assigns a higher rank to the target enzymes compared to CLIPZyme.

| Enzyme/Reaction/EC | Method | Reaction Prediction Rank |
|---|---|---|
| A0A0N0U829/<br>RHEA:51608/<br>1.1.99.40 | *Horizyn-1* | 4 |
| | MMSeqs2 | 19 |
| | BLASTp | 19 |
| | Foldseek | Unranked |
| | CLEAN | EC Not Predicted |
| | CLIPZyme | 26 |
| A0A0N0BLM2/<br>RHEA:25094/<br>2.7.4.14,2.7.4.25 | *Horizyn-1* | 1 |
| | MMSeqs2 | Unranked |
| | BLASTp | Unranked |
| | Foldseek | Unranked |
| | CLEAN | EC Not Predicted |
| | CLIPZyme | 624 |

TABLE S7. **Comparison of *Horizyn-1* and standard annotation tools on protein function prediction of *Thermus aquaticus* enzymes.** The predicted rank (or success) of the target reactions (see Fig. 2) is listed with respective screening sets. *Horizyn-1* predicted the newly confirmed *(R)*-2-hydroxyglutarate–pyruvate transhydrogenase activity highly at 4, while MMSeqs2 and BLASTp ranked it 19, and Foldseek and CLEAN failed to rank/predict it at all. Only *Horizyn-1* predicted the dCMP kinase activity of A0A0N0BLM2, while no other method managed to rank/predict the activity at all (see Protein Function Annotation Comparison). "Unranked" indicates the method failed to identify any sequence with the desired function as a positive hit. The CLEAN webserver predicts a single EC number; "Predicted"/"Not Predicted" indicates whether the predicted EC number matched that of the target reaction.

| Reaction | Enzyme | | | | | | | |
|---|---|---|---|---|---|---|---|---|
| | Q9HUI9 (TA1) | | P09053 (TA2) | | Q9I6J2 (TA3) | | O50131 (TA4) | |
| | *Horizyn-1* | CLIPZyme | *Horizyn-1* | CLIPZyme | *Horizyn-1* | CLIPZyme | *Horizyn-1* | CLIPZyme |
| PG-$\alpha$ | 14 | 920 | 2 | 2318 | 48 | 5712 | 2897 | N/A |
| PG-$\varepsilon$ | 109 | 931 | 35 | 1322 | 6 | 1558 | 15 | N/A |
| 2-TPG-$\alpha$ | 20 | 1333 | 13 | 3229 | 18 | 4435 | 2120 | N/A |
| 2-TPG-$\varepsilon$ | 1061 | 1240 | 949 | 1247 | 77 | 1513 | 1 | N/A |
| 2-FG-$\alpha$ | 7 | 953 | 8 | 2622 | 47 | 5433 | 2826 | N/A |
| 2-FG-$\varepsilon$ | 1115 | 1224 | 366 | 1273 | 109 | 1452 | 4 | N/A |
| Max Rank | 7 | 931 | 2 | 1247 | 6 | 1452 | 1 | N/A |

TABLE S8. **Comparison of *Horizyn-1* and CLIPZyme on enzyme discovery for non-canonical amino acid synthesis.** The predicted rank of each target enzyme (see Fig. 3) is listed within each model's respective screening set. *Horizyn-1*'s screening set consists of ∼7 million enzymes, while CLIPZyme considers ∼260,000. The last row lists the best (lowest) rank achieved by each enzyme across all transaminase reactions. Despite a substantially larger screening set, *Horizyn-1* consistently assigns a higher rank (top 10) to the target enzymes compared to CLIPZyme (ranks 900–1500).

| Enzyme Class | SMARTS Template |
|---|---|
| Aminotransferases | [CH:1]([NH2:3])[C:2](=[O:4])[OH:5].[CH0:6](=[O:8])[C:7](=[O:9])[OH:10]>><br>[CH0:1](=[O:8])[C:2](=[O:4])[OH:5].[C@H:6]([NH2:3])[C:7](=[O:9])[OH:10] |
| | [CH2:1]([NH2:3])[C:2](=[O:4])[OH:5].[CH0:6](=[O:8])[C:7](=[O:9])[OH:10]>><br>[CH:1](=[O:8])[C:2](=[O:4])[OH:5].[C@H:6]([NH2:3])[C:7](=[O:9])[OH:10] |
| DUF849 | [C:1](=[O:2])[CH2:3][C:4](=[O:5])[OH:6].[CH3:7][C:8](=[O:9])[S:10][CH2:11]>><br>[CH3:7][C:8](=[O:9])[CH2:3][C:4](=[O:5])[OH:6].[C:1](=[O:2])[S:10][CH2:11] |
| Esterases | [C,c:1][OH0:2][C:3]=[O:4].[OH2:5]>>[C,c:1][OH:2].[C:3](=[O:4])[OH:5] |
| Nitrilases | [C,c:1][C:2]#[N:3].[OH2:4].[OH2:5]>>[C,c:1][C:2](=[OH0:4])([OH:5]).[NH3:6] |
| OleA Thiolases | [O:4]=[CH0:3]([OH0:5][C,c:6])[C,c:7].[CH2:1][SH:2]>><br>[CH2:1][SH0:2][CH0:3](=[O:4])[C,c:7].[OH:5][C,c:6] |
| Phosphatases | [O:4]=[P:3]([OH:6])([OH:5])[OH0:2][*:1].[OH2:7]>>[OH:2][*:1].[OH:7][P:3]([OH:6])([OH:5])=[O:4] |
| | [O:4]=[P:3]([OH:6])([OH:5])[NH:2][*:1].[OH2:7]>>[NH2:2][*:1].[OH:7][P:3]([OH:6])([OH:5])=[O:4] |
| | [O:4]=[P:3]([OH:6])([OH:5])[NH2:1].[OH2:7]>>[NH3:1].[OH:7][P:3]([OH:6])([OH:5])=[O:4] |
| | [O:4]=[P:3]([OH:6])([OH:5])[C:2][*:1].[OH2:7]>>[O:4]=[P:3]([OH:6])([OH:5])[OH:7].[C:2][*:1] |
| | [O:4]=[P:3]([OH0:6])([OH:5])[OH0:2][P:1].[OH2:7]>>[O:4]=[P:3]([OH0:6])([OH:5])[OH:2].[OH:7][P:1] |

TABLE S9. R**eaction SMARTS templates for reactions in the enzyme promiscuity benchmark.** See Enzyme Promiscuity Benchmark.

| Enzyme Class | Active Substrates | Inactive Substrates |
|---|---|---|
| Aminotransferases | 186 | 264 |
| DUF849 | 274 | 2463 |
| Esterases | 3076 | 10940 |
| Nitrilases | 85 | 599 |
| OleA Thiolases | 550 | 545 |
| Phosphatases | 5357 | 30177 |

TABLE S10. **Counts of active and inactive enzyme-substrate pairs in enzyme promiscuity benchmark.** See Enzyme Promiscuity Benchmark.

| Analyte | Ion Mode | MRM Transitions ($m/z$) | Retention Time (min) |
|---|---|---|---|
| 2-furylglycine | Positive | *142.2 → 97.1*<br>142.2 → 69.2 | 2.2 |
| L-phenylglycine | Positive | 152.2 → 135.0<br>*152.2 → 107.2*<br>152.2 → 79.2 | 3.0 |
| 2-thienylglycine | Positive | 158.1 → 141.2<br>*158.1 → 113.1*<br>158.1 → 85.1 | 2.7 |
| CDP | Negative | *402.1 → 158.7*<br>402.1 → 110.1<br>402.1 → 78.9 | 2.4 |
| dCDP | Negative | *386.0 → 158.7*<br>386.0 → 78.7 | 2.4 |

TABLE S11. **Multiple reaction monitoring (MRM) $m/z$ transitions.** Ion mode, $m/z$ transitions, and retention times for analytes measured in this study. Transitions in italics represent data used for analysis.

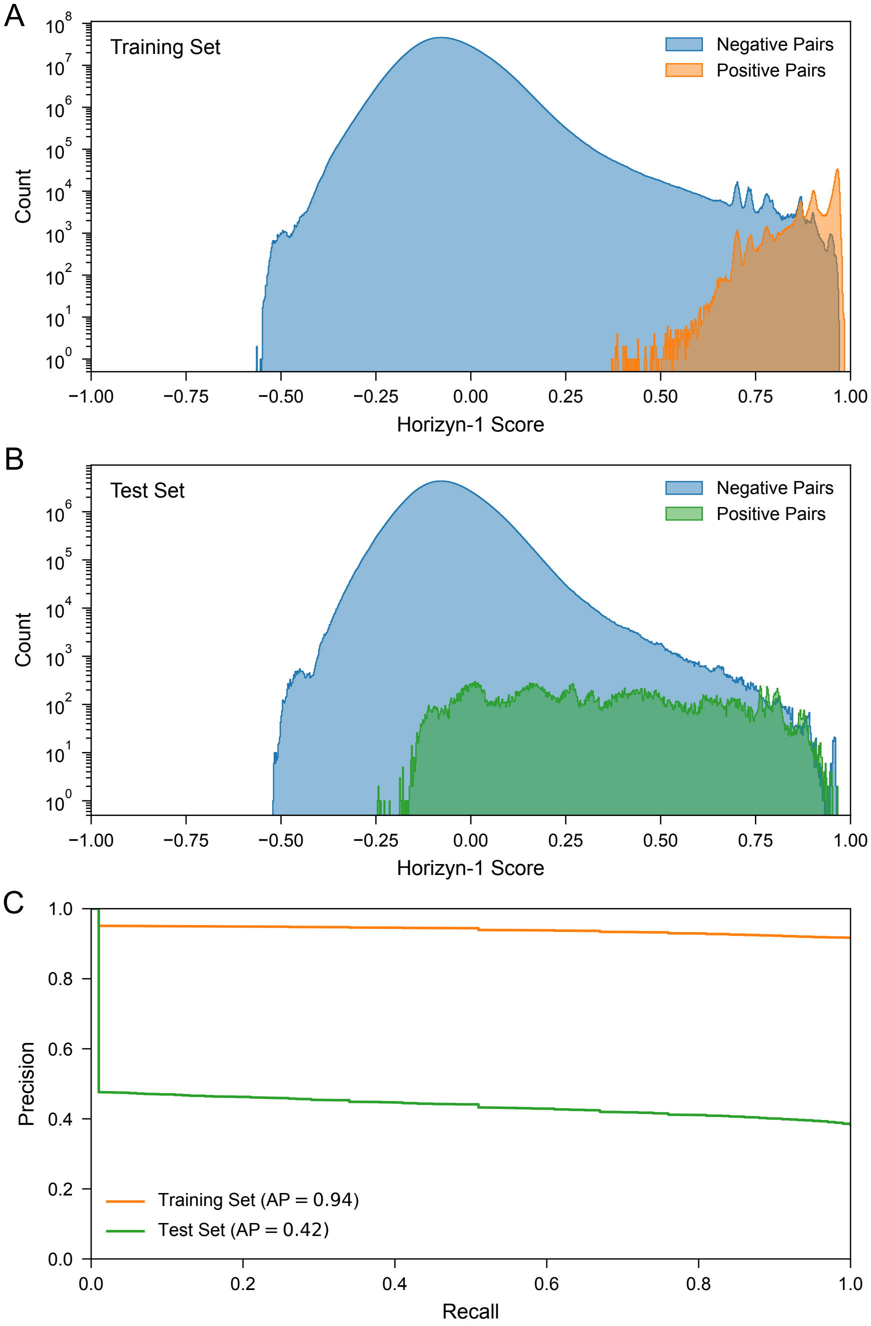


FIG. S1. **Score distributions and precision-recall curves.** **(A)** Score distributions for positive (orange) and negative (blue) pairs in the development model training set. **(B)** Score distributions for positive (green) and negative (blue) pairs in the development model test set. **(C)** Mean precision-recall curves for classifying positive versus negative enzymes in the development model training set (orange) and test set (green). The mean average precision (area under the precision-recall curve) is 0.94 for the training set and 0.42 for the test set. The mean curve was calculated by averaging precision values at each recall threshold across individual reactions.

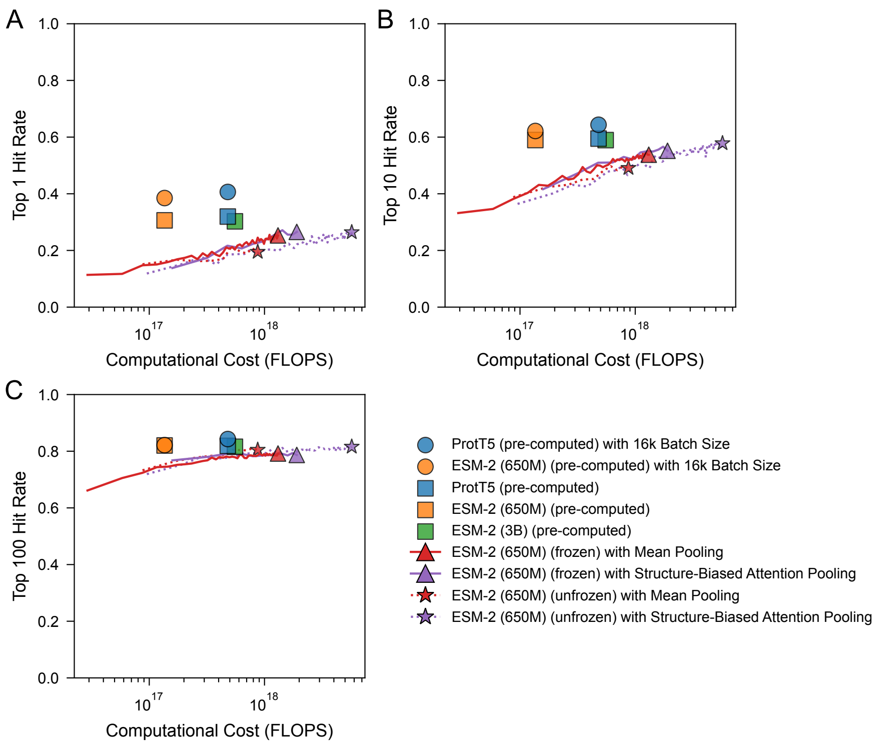


FIG. S2. **Performance vs. computational cost for different protein embedding strategies.** The trade-off between model performance and computational efficiency is shown for various protein language model (pLM) embedding strategies. The **(A)** top-1, **(B)** top-10, and **(C)** top-100 hit rates on the validation set are plotted against the cumulative computational cost, measured in floating point operations (FLOPs). Lines represent the performance trajectory during training, while the solid markers indicate the final performance of each fully trained model. Two main paradigms are compared: models trained on pre-computed pLM embeddings (circles and squares) and models that integrate a live ESM-2 (650M) model during training (triangles and stars). The live models were trained with either a frozen or unfrozen pLM, and with or without a structure-aware attention pooling layer. See Protein Language Model Compute Cost and Finetuning Analysis for details.

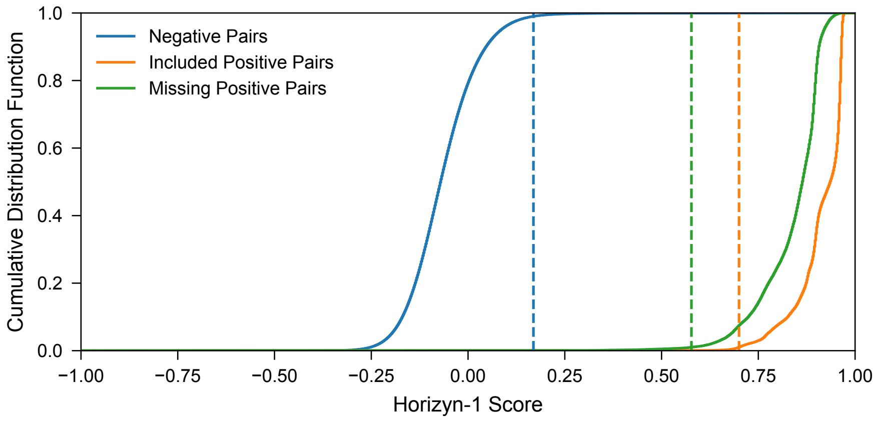


FIG. S3. **Performance on unannotated positive reaction-enzyme pairs.** Cumulative distribution functions (CDFs) of model scores for negative (blue), included positive (orange), and missing positive (green) pairs. Vertical lines mark the 99th percentile for negative pairs (0.17) and the 1st percentile for included (0.70) and missing (0.58) positive pairs. The negative and included thresholds are identical to the baseline model. Notably, performance on missing pairs remains high, with a threshold (0.58) comparable to included pairs (0.70). Furthermore, 93% of missing pairs scoring above the included pair threshold in comparison to only 0.1% of negative pairs. See Missing Pair Analysis for more details..

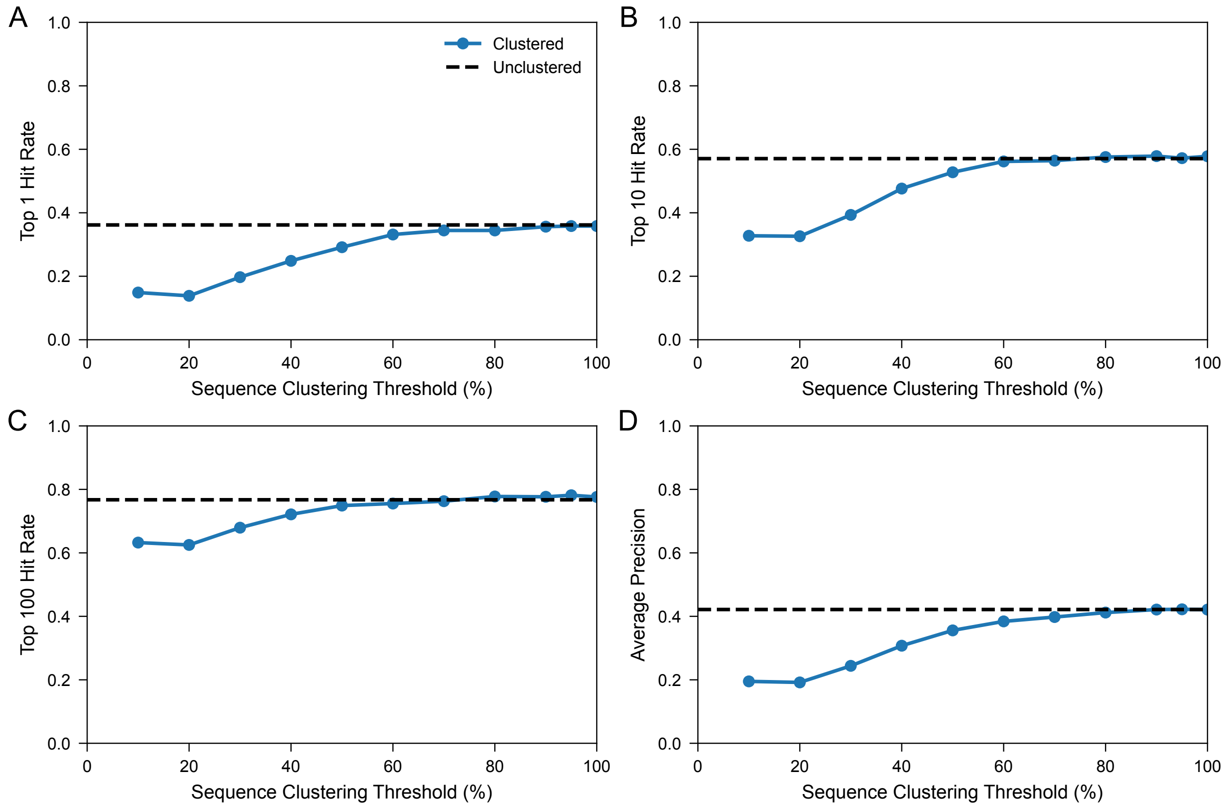


FIG. S4. **Performance vs. sequence identity clustering threshold.** The **(A)** top-1 hit rate, **(b)** top-10 hit rate, **(C)** top-100 hit rate, and **(D)** average precision are shown for the development model on the test set for varying levels of clustering on the protein sequences in the training set (see Sequence Clustering Analysis for details). Metrics for the unclustered training set are shown for comparison (see Fig. 1). A sequence identity of threshold of 80% was ultimately chosen for the dataset used to train the inference model (see Inference Dataset) to compress the size of the dataset as much as possible without a significant drop in performance.

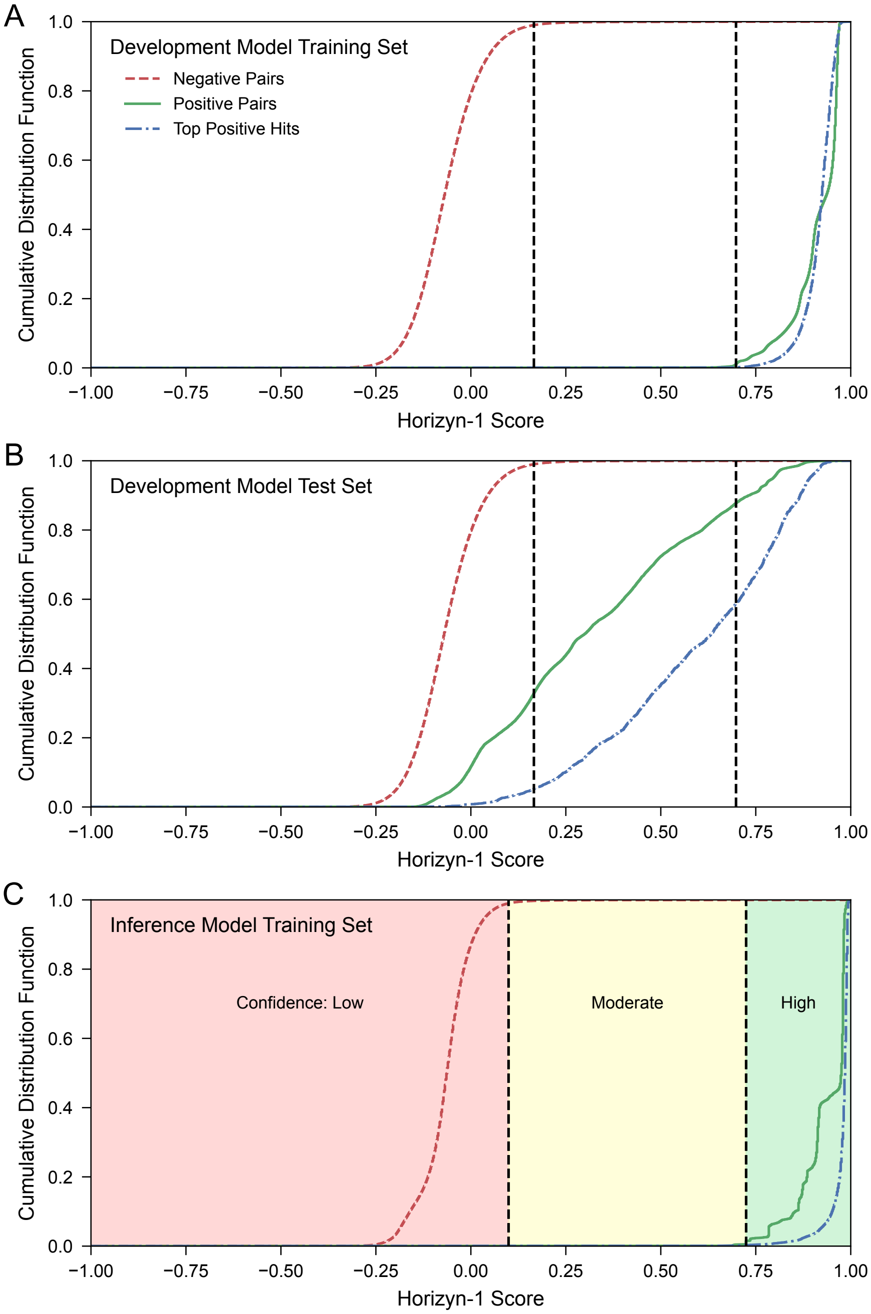


FIG. S5. **Score cumulative distribution functions and confidence heuristic.** Cumulative Distribution Functions (CDFs) of model scores are shown for three categories: negative pairs (red), positive pairs (green), and the top positive hit for each reaction (blue). Score distributions generated by the model trained on the training split of the development dataset are plotted for **(A)** the development training set and **(B)** the development model test set, demonstrating clear separation between the scores of positive and negative pairs. **(C)** The score distributions generated by *Horizyn-1* for the inference training set are used to define a three-tiered confidence heuristic, visually represented by the shaded regions. The dashed lines denote boundaries between "Low," "Moderate," and "High" confidence predictions, which are based on percentiles of the negative and positive score distributions (see Model Confidence Heuristic).

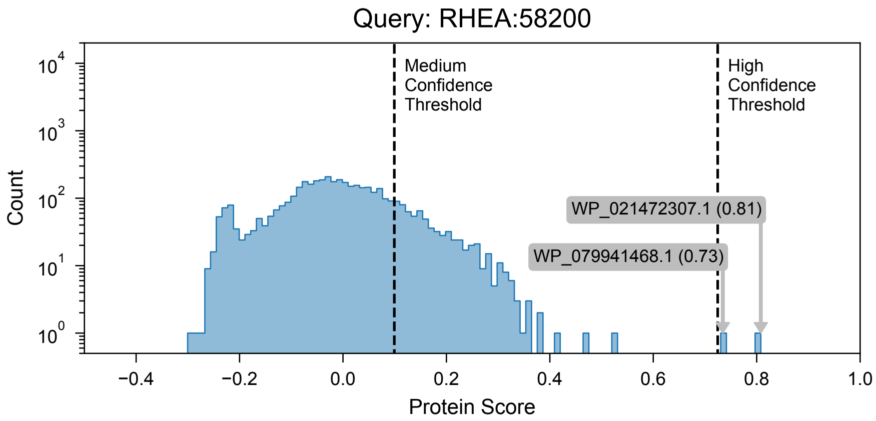


FIG. S6. ***Horizyn-1* predictions for 6-aminohexanoate transaminase orphan reaction (RHEA:58200) among *Arthrobacter* sp. KI72 genome.** Distribution of *Horizyn-1* protein scores for the 6-aminohexanoate aminotransferase orphan reaction (RHEA:58200) queried against all 4,173 protein-coding sequences from the *Arthrobacter* sp. KI72 genome. The scores for the two highest-ranking proteins, WP_021472307.1 (a GabT homolog) and WP_079941468.1, are explicitly annotated (0.81 and 0.73, respectively). Both candidates surpass the high-confidence threshold (right dashed line), supporting their selection for subsequent experimental characterization.

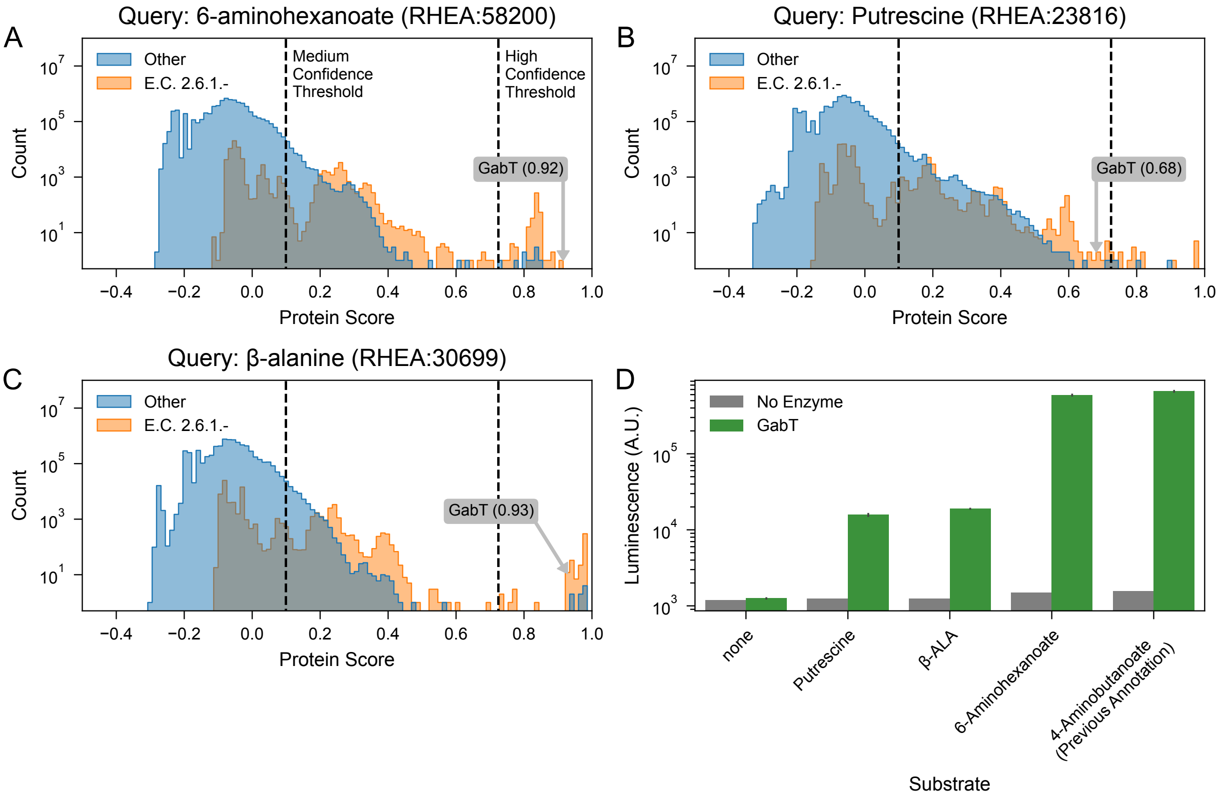


FIG. S7. ***Horizyn-1* prediction and experimental confirmation of promiscuous $\omega$-transaminase activity for *E. coli* GabT. (A–C)** Distributions of *Horizyn-1* protein scores for three orphan $\omega$-transaminase reactions queried against our inference screening set of ~7 million enzymes. The reactions shown are for **(A)** 6-aminohexanoate (RHEA:58200), **(B)** putrescine (RHEA:23816), and **(C)** $\beta$-alanine (RHEA:30699). In each plot, the score for *E. coli* GabT (P22256) is highlighted with its score in parentheses, ranking it as a high-confidence candidate for these promiscuous activities. The histograms differentiate between known transaminases (EC 2.6.1.-; orange) and all other enzymes (blue). **(D)** Experimental validation of GabT activity. L-glutamate production (measured as luminescence on a logarithmic scale) was assayed for the predicted substrates and the previously annotated substrate, 4-aminobutanoate (RHEA:23352). Green bars indicate activity with purified GabT, while gray bars represent the background signal from the uncatalyzed reaction. The results confirm that GabT has strong activity with 6-aminohexanoate and displays weaker, promiscuous activity with putrescine and $\beta$-alanine. Error bars denote the standard deviation from duplicate measurements.

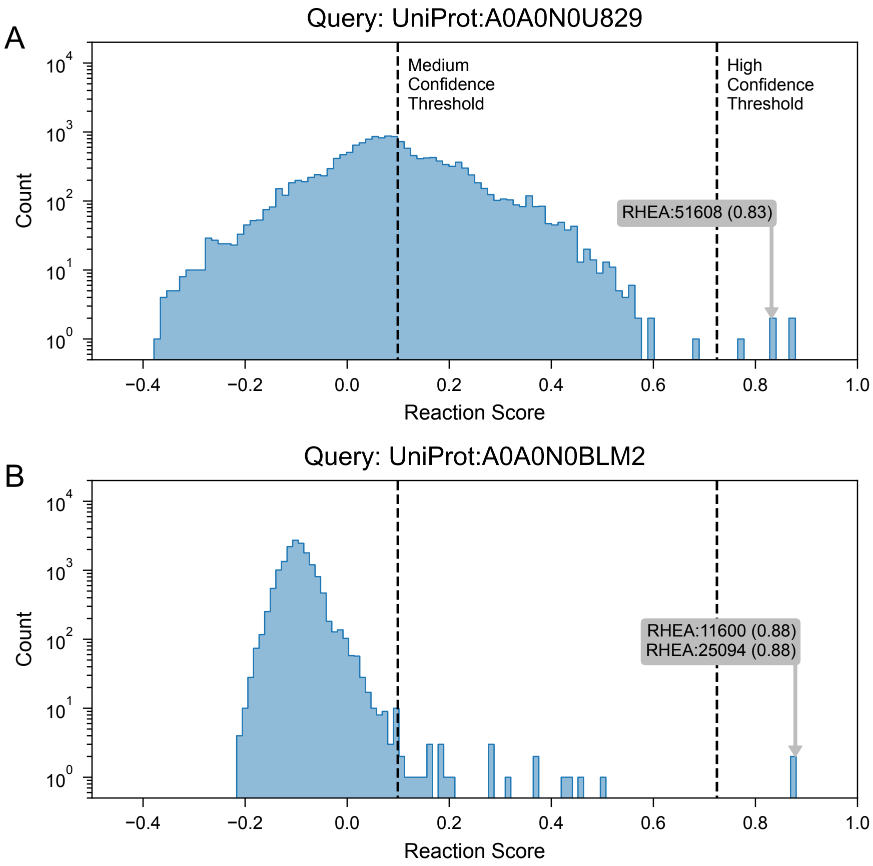


FIG. S8. ***Horizyn-1* predictions for promiscuous activity for enzymes from *Thermus aquaticus*. (A)** Distribution of *Horizyn-1* reaction scores for the hypothetical D-lactate dehydrogenase A0A0N0U829, queried against all 15,814 Rhea reactions. While the protein's annotated function is lactate oxidation, the analysis predicted a high-scoring promiscuous activity: *(R)*-2-hydroxyglutarate–pyruvate transhydrogenase (RHEA:51608, score = 0.83). This prediction, which surpassed the high-confidence threshold, suggested the enzyme could utilize alternative electron donor/acceptor pairs and was subsequently confirmed experimentally. **(B)** Distribution of *Horizyn-1* reaction scores for the uncharacterized protein A0A0N0BLM2, queried against all Rhea reactions. The model predicted two high-scoring kinase activities: cytidine monophosphate (CMP) kinase (RHEA:11600, score = 0.88) and deoxycytidine monophosphate (dCMP) kinase (RHEA:25094, score = 0.88). Subsequent experimental testing confirmed the dCMP kinase activity, demonstrating *Horizyn-1*'s ability to successfully assign function to a previously unannotated protein.

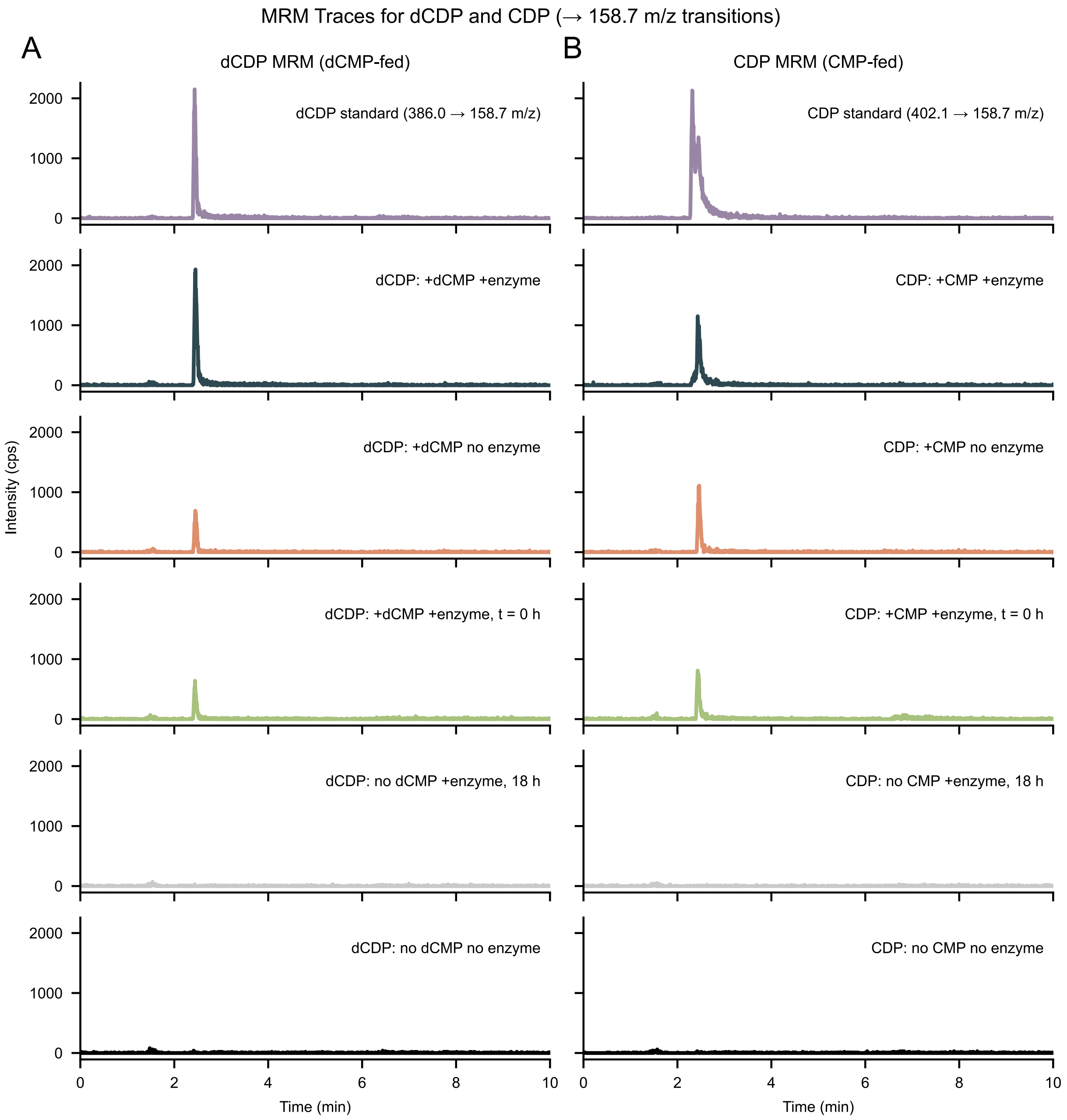


FIG. S9. **MRM analysis of (d)CMP kinase activity on dCMP and CMP substrates.** Multiple reaction monitoring (MRM) chromatograms show extracted ion transitions for **(A)** dCDP (386.0 → 158.7 $m/z$, left) and **(B)** CDP (402.1 → 158.7 $m/z$, right) under various reaction conditions. Each row corresponds to a unique condition: reactions with or without enzyme (A0A0N0BLM2), with or without nucleotide substrate (dCMP or CMP), and an early timepoint control ($t$ = 0 h). A strong dCDP signal is observed only in the presence of both dCMP and enzyme, indicating conversion of dCMP to dCDP after 18 h of incubation. In contrast, no appreciable CDP signal is observed relative to controls in reactions containing CMP and enzyme. These results demonstrate that the enzyme selectively phosphorylates dCMP but not CMP under the conditions tested.

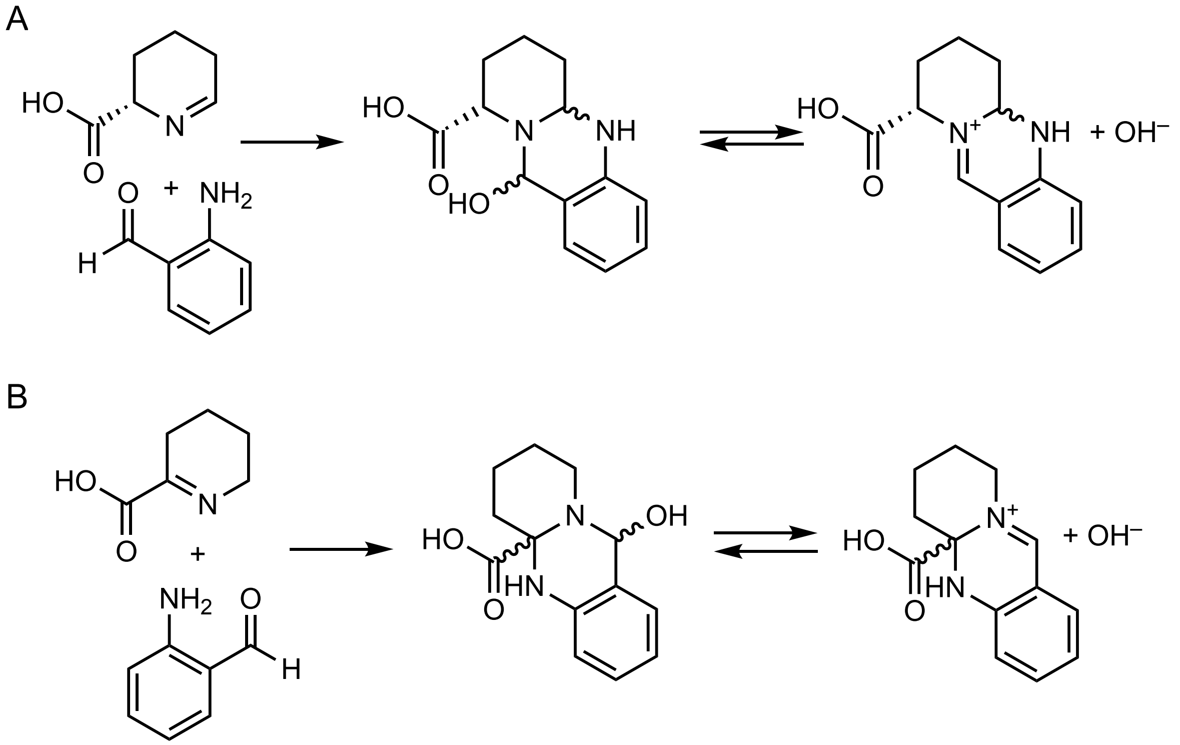


FIG. S10. **Reaction schemes for l-lysine-driven transamination assay.** Reaction schemes for the detection of lysine **(A)** $\varepsilon$-amine and **(B)** $\alpha$-amine transamination products via *o*-aminobenzaldehyde reactivity. Lysine transaminase screening may be performed colorimetrically by taking advantage of the spontaneous cyclization of products that react with *o*-aminobenzaldehyde. Transamination utilizing the $\varepsilon$-amine of lysine produces L-allysine which spontaneously cyclizes to form *(S)*-1-piperideine-6-carboxylate. This molecule reacts with *o*-aminobenzaldehyde to form a three-membered ring product with an absorbance maximum at 465 nm [4, 6, 10]. Quantification can proceed using the extinction coefficient of 2.8 $mM^{-1}cm^{-1}$. One unit of enzyme activity is defined as catalyzing formation of 1 nmol of product per min. Similarly, transamination utilizing the $\alpha$-amine of lysine produces 6-amino-2-oxohexanoic acid which spontaneously cyclizes to form 1-piperideine-2-carboxylate. Reaction with *o*-aminobenzaldehyde produces a three-membered ring product with an absorbance maximum of 450 nm [4, 6, 10].

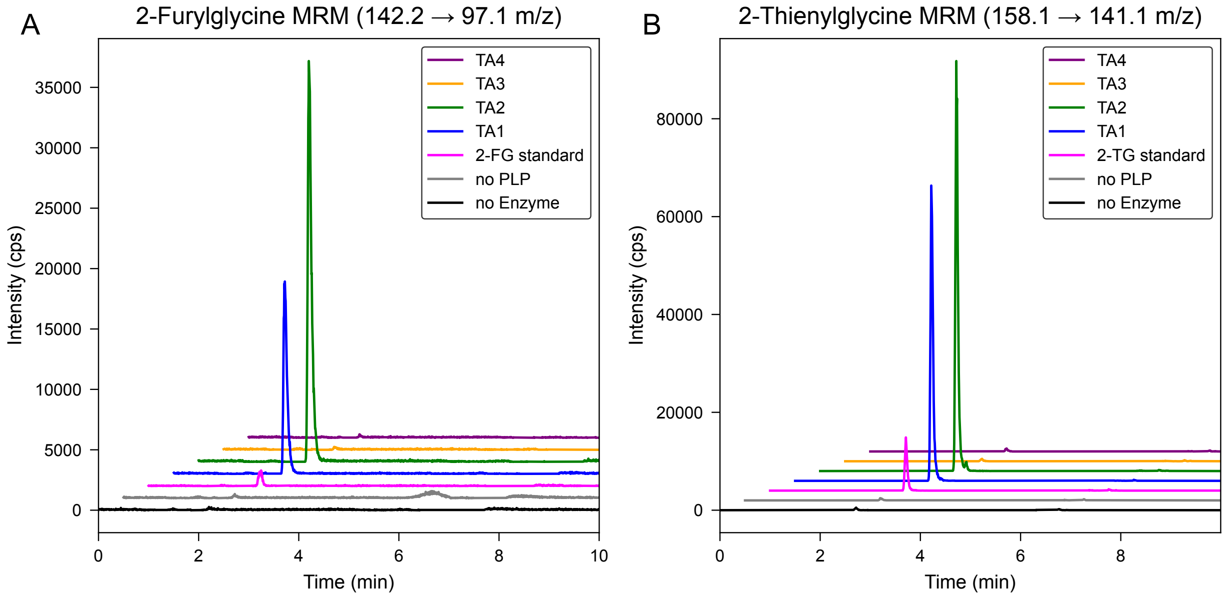


FIG. S11. **MRM for non-canonical amino acid detection.** Positive mode MRM of characteristic transitions for non-canonical amino acids synthesized via smart lysine transaminases. Traces are shown for **(A)** 2-furylglycine and **(B)** 2-thienylglycine. Negative controls in which the PLP cofactor or the enzyme were omitted from the reaction are shown in gray and black traces, respectively.

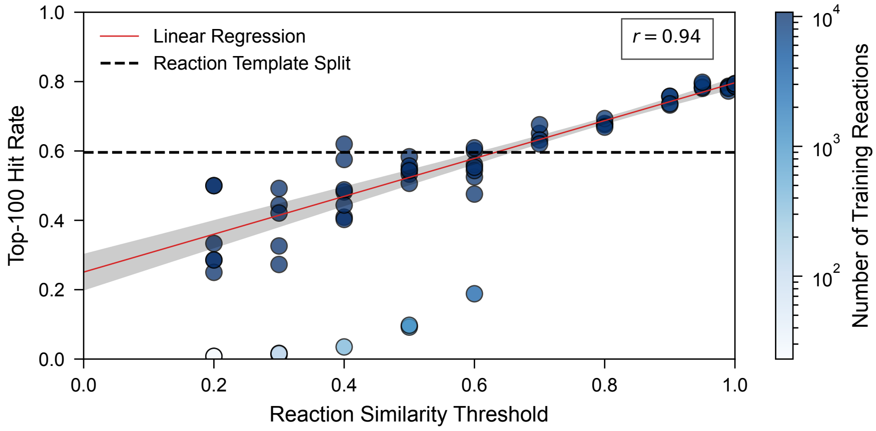


FIG. S12. **Impact of training and validation data similarity on model performance.** The scatter plot shows the top-100 hit rate for models trained on data splits generated using various fingerprint-based reaction similarity thresholds. A lower threshold creates a more challenging split by increasing the dissimilarity between reactions in the training and validation sets. Each point represents an independently trained model, colored by the number of training reactions in its respective split (color bar). A linear regression (red line) with 95% confidence intervals (gray shading) demonstrates a clear positive correlation between the similarity threshold and model performance (Pearson's $r = 0.94$). For comparison, the dashed line shows the average performance (top-100 hit rate of 60%) when splitting data by reaction templates (see Reaction Similarity Clustering Analysis).

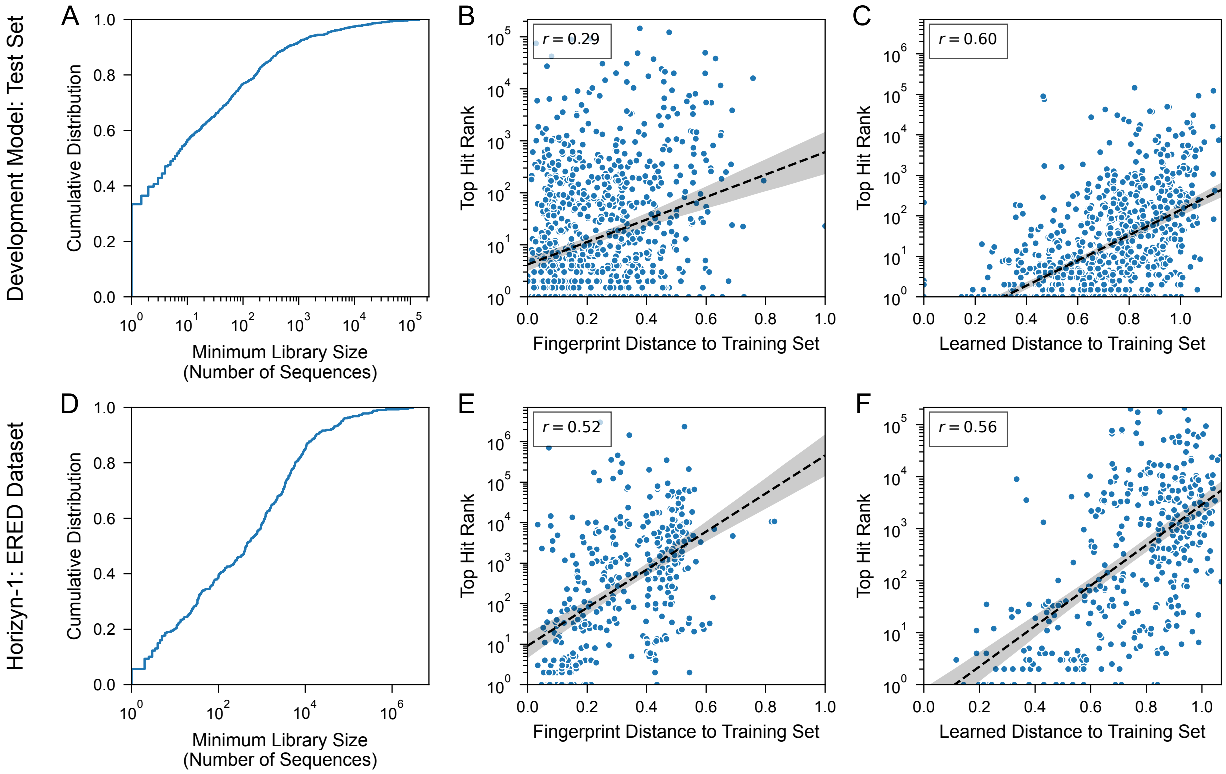


FIG. S13. **Model performance and reaction novelty.** Model performance and its relationship to reaction novelty are presented for **(A–C)** a model trained on the development dataset training split and evaluated on the development test split and **(D–F)** *Horizyn-1* (trained on the inference dataset) evaluated on the ERED dataset. **(A, D)** The cumulative distribution of the top hit ranks; the $x$-axis represents the minimum number of sequences that would need to be experimentally screened to find the first positive hit for a given reaction. For a given position $k$ on the $x$-axis, the position on the $y$-axis is the top-$k$ hit rate. **(B, C, E, F)** Scatter plots correlating the rank of the top-scoring enzyme hit with the reaction's novelty. Novelty is defined as the minimum distance from a test reaction to any reaction in the corresponding training set. This distance is measured using either **(B, E)** Tanimoto distance of chemical fingerprints (see Structural Fingerprint Clustering Protocol) or **(C, F)** Euclidean distance within the learned embedding space. For both datasets, the performance is more strongly correlated with the learned distance than the fingerprint distance (Pearson's $r = 0.60$ vs. 0.29 for the development test set; $r = 0.56$ vs. 0.52 for the ERED set), demonstrating that the model's learned embedding space is a more effective measure of biochemical novelty.

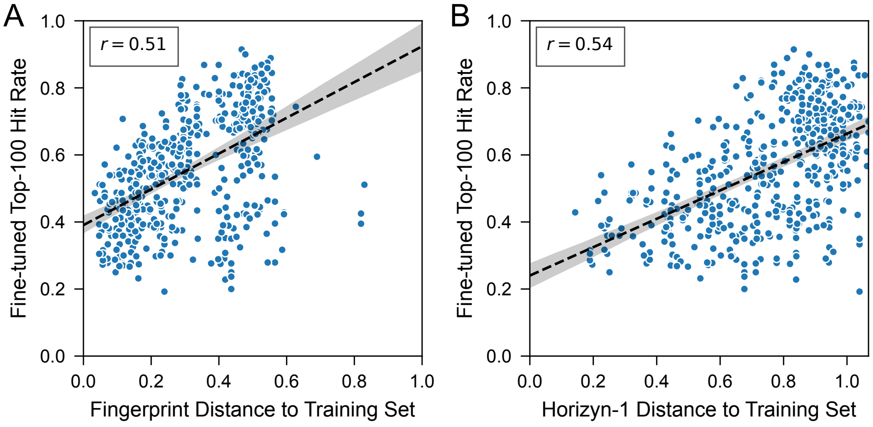


FIG. S14. **Fine-tuning improvement vs. reaction novelty.** The scatter plots correlate the top-100 hit rate on a random ERED validation set after fine-tuning the *Horizyn-1* inference model on a single reaction with the novelty of that reaction (see Ene-Reductase Fine-tuning Protocol). Novelty is quantified as the minimum distance of the reaction used for fine-tuning to any reaction in the original training set, measured using either **(A)** chemical fingerprint Tanimoto distance (see Structural Fingerprint Clustering Protocol) or **(B)** Euclidean distance within the *Horizyn-1* embedding space. The performance after fine-tuning is strongly correlated with both distances ($r = 0.54$ vs. 0.51), suggesting that biochemical novelty leads to performance improvements upon fine-tuning.

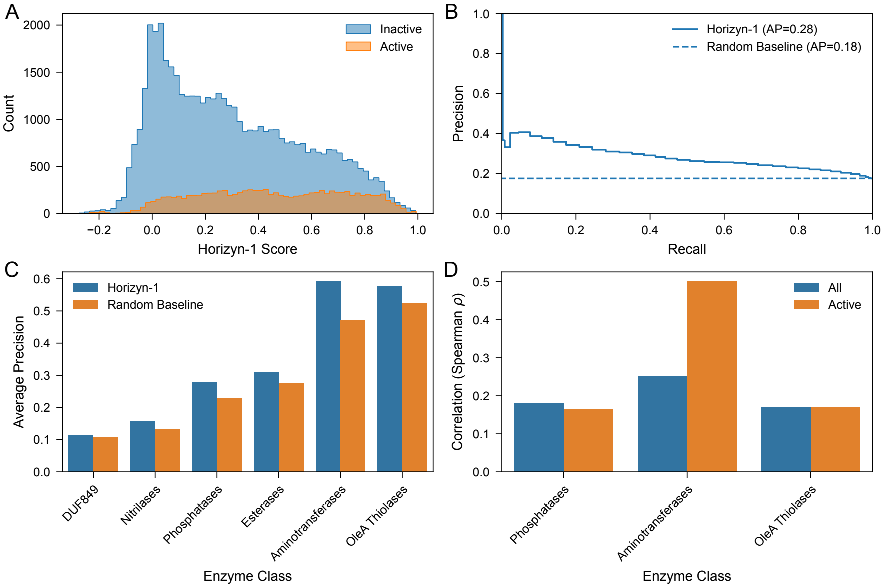


FIG. S15. **Performance on positive/negative activity prediction.** *Horizyn-1*'s ability to distinguish between reactions with active and inactive substrates for specific classes of enzymes was evaluated using standard benchmark dataset for enzyme promiscuity (`https://github.com/samgoldman97/enzyme-datasets`). **(A)** Score distributions for inactive (blue) an active (orange) substations. **(B)** Precision-recall curve for classifying the active versus inactive substrates compared to a random baseline. **(C)** Average precision of *Horizyn-1* and a random baseline for each enzyme class. **(D)** Spearman rank correlation coefficients between *Horizyn-1* scores and activity across available enzyme classes for all enzyme-substrate pairs (blue) and active pairs only (orange). $p$-values for all correlation coefficients were less than $10^{-4}$.
